# Incomplete proteasomal degradation of green fluorescent proteins in the context of tandem fluorescent protein timers

**DOI:** 10.1101/023119

**Authors:** Anton Khmelinskii, Matthias Meurer, Chi-Ting Ho, Birgit Besenbeck, Julia Füller, Marius K. Lemberg, Bernd Bukau, Axel Mogk, Michael Knop

## Abstract

Tandem fluorescent protein timers (tFTs) report on protein age through time-dependent change in color, which can be exploited to study protein turnover and trafficking. Each tFT, composed of two fluorescent proteins (FPs) that differ in maturation kinetics, is suited to follow protein dynamics within a specific time range determined by the maturation rates of both FPs. So far tFTs were constructed by combining different slower-maturing red fluorescent proteins (redFPs) with the same faster-maturing superfolder green fluorescent protein (sfGFP). Towards a comprehensive characterization of tFTs, we compare here tFTs composed of different faster-maturing greenFPs, while keeping the slower-maturing redFP constant (mCherry). Our results indicate that the greenFP maturation kinetics influences the time range of a tFT. Moreover, we observe that commonly used greenFPs can partially withstand proteasomal degradation due to the stability of the FP fold, which results in accumulation of tFT fragments in the cell. Depending on the order of FPs in the timer, incomplete proteasomal degradation either shifts the time range of the tFT towards slower time scales or precludes its use for measurements of protein turnover. We identify greenFPs that are efficiently degraded by the proteasome and provide simple guidelines for design of new tFTs.

**Abbreviations:** tFT – tandem fluorescent protein timer

FP – fluorescent protein

greenFP – green fluorescent protein

redFP – red fluorescent protein

## Introduction

Fluorescent proteins (FPs) become fluorescent only upon correct folding of the polypeptide and formation of the fluorophore through a series of chemical reactions within the β-barrel fold. This maturation process takes place on time scales from minutes to hours, depending on the FP and the environment (Tsien, 1998; Shaner *et al.*, 2005). The time delay between protein synthesis and completion of the maturation process is exploited in tandem fluorescent protein timers (tFTs) to measure the dynamics of various cellular processes (Khmelinskii *et al.*, 2012). A tFT is a fusion of two FPs that differ in maturation kinetics and spectral properties (i.e. color), e.g. a slower-maturing red fluorescent protein (redFP) and a faster-maturing green fluorescent protein (greenFP) (Figure 1A). The greenFP is typically selected to be as fast-maturing as possible and thus reports on protein localization and abundance. Because of the difference in maturation kinetics between the greenFP and redFP moieties, the color of a tFT (defined as the redFP/greenFP ratio of fluorescence intensities) changes over time (Figure 1A). When a tFT is fused to a protein of interest, its color provides a measure of protein age that can be used to follow protein degradation, trafficking and segregation during cell division (Khmelinskii *et al.*, 2012).

**Figure 1.**
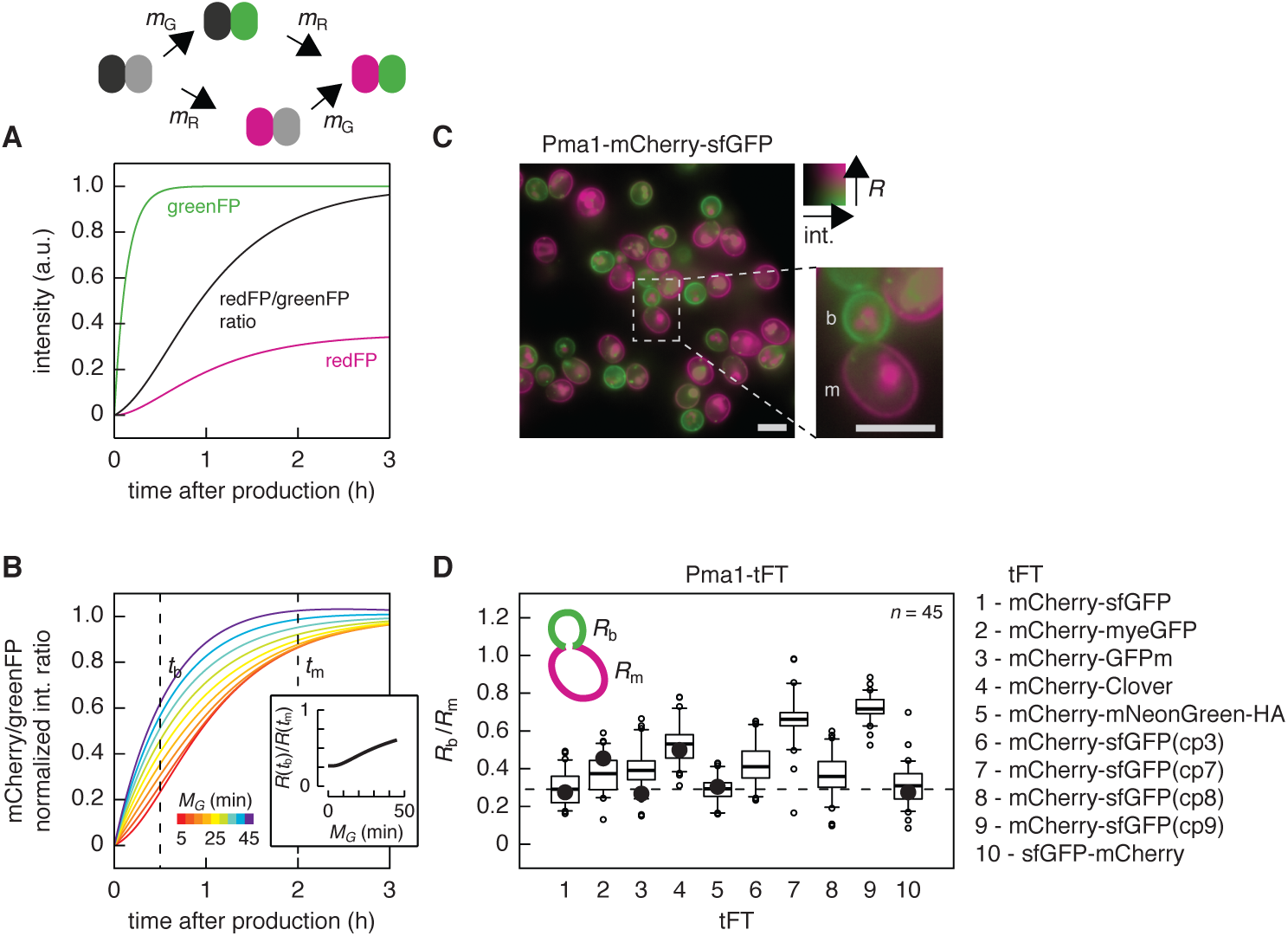
Analysis of protein age with different tFTs. A – Behavior of a tFT composed of a slower-maturing redFP (black – magenta, maturation rate constant *m_R_*) and a faster-maturing greenFP (grey – green, maturation rate constant *m_G_*). Fluorescence intensity curves were calculated using published maturation parameters for mCherry (redFP) and sfGFP (greenFP) (Khmelinskii *et al.*, 2012) for a population of tFT molecules initialized in the non-mature state in the absence of protein production and degradation. Intensity curves are normalized to the brightness of sfGFP, ratio curve is normalized to the point of complete maturation. B – tFTs composed of mCherry and greenFPs with different maturation kinetics. mCherry/greenFP intensity ratio curves were calculated as in A using published maturation parameters for mCherry (Khmelinskii *et al.*, 2012) and a maturation half-time *M*_G_ between 5 and 45 min for greenFP. Note that the maturation half-time *M*_G_ is related to the maturation rate constant *m*_G_ as *M*_G_ = ln(2)/*m*_G_. Each curve is normalized to the point of complete maturation. Curves are color-coded according to *M*_G_ as indicated. Inset shows the comparison of mCherry/greenFP intensity ratios (R) at two time points – *t*_b_ = 30 min (approximate age of a bud) and *t*_m_ = 2 h (approximate age of a young mother cell) – as a function of *M*_G_. C – Representative intensity-weighted ratiometric image of cells expressing Pma1-mCherry-sfGFP, color-coded according to sfGFP intensity (int.) and mCherry/sfGFP intensity ratio (*R*). A dividing cell with labeled mother (m) and bud (b) compartments is shown. Scale bars are 5 μm. D – mCherry/greenFP intensity ratios (*R*) of Pma1 tagged with the indicated tFTs were measured at the plasma membrane in pairs of mother (*R*_m_) and bud (*R*_b_) cells (*n* = 45 pairs for each tFT). Center lines mark the medians, box limits indicate the 25th and 75th percentiles, whiskers extend to 5th and 95th percentiles. The difference between *R*_m_ and *R*_b_ is significant for all tFTs (p < 10^−19^ in a paired t-test). The mCherry-mNeonGreen timer carried a C-terminal hemagglutinin epitope (HA) tag to facilitate detection by immunoblotting. Black circles mark theoretical *R*_b_/*R*_m_ values calculated according to a simple model of mCherry-greenFP maturation in B using 30 min and 2 h as approximate ages of a bud and a young mother cell, respectively, and published maturation half-times for mCherry and different greenFPs (Khmelinskii *et al.*, 2012).

The range of protein ages that can be analyzed with a tFT, i.e. its time range, is strongly influenced by the maturation kinetics of the slower-maturing FP in the pair (Khmelinskii *et al.*, 2012). Therefore, in the tFTs described thus far (Khmelinskii *et al.*, 2012; Dona *et al.*, 2013; Khmelinskii and Knop, 2014), different slower-maturing redFPs (mCherry, TagRFP, DsRed1) were combined with the same faster-maturing superfolder green fluorescent protein (sfGFP) (Pédelacq *et al.*, 2006). Several properties make sfGFP a greenFP of choice for construction of tFTs: it is bright, it folds independently of the protein it is fused to, it is one of the faster-maturing greenFPs, with a maturation half-time of ~6 min in yeast (Pédelacq *et al.*, 2006; Khmelinskii *et al.*, 2012), and an additional V206R mutation ensures it is monomeric (Zacharias *et al.*, 2002). Nevertheless, there is a wide range of greenFPs with other desirable properties such as high molecular brightness or high photostability (Dean and Palmer, 2014). However, their *in vivo* maturation rates, which determine their suitability for construction of new tFTs (Figure 1B), are not well characterized.

Here we sought to compare the performance of different greenFPs within tFTs. We constructed a series of mCherry-greenFP timers and examined their suitability for comparative measurements of protein age and protein degradation kinetics in yeast. We identified the stability of the greenFP fold as a new parameter that affects tFT behavior and directs the design and use of tFT as reporters of protein degradation.

## Results

We constructed new putative tFTs by fusing the slower-maturing red fluorescent protein mCherry (Shaner *et al.*, 2004) to different greenFPs: the commonly used monomeric yeast codon-optimized enhanced GFP (myeGFP) (GFPmut 3a from (Cormack *et al.*, 1996) with the V206R dimerization-preventing mutation (Zacharias *et al.*, 2002)), the fast-maturing GFPm (Yoo *et al.*, 2007; Iizuka *et al.*, 2011), and two greenFPs with superior brightness – Clover (Lam *et al.*, 2012) and mNeonGreen (Shaner *et al.*, 2013) (Figure S1, A and B). myeGFP, GFPm, Clover and mNeonGreen have red-shifted excitation peaks compared to sfGFP. Using such greenFPs is likely to improve signal-to-noise ratio in fluorescence imaging as autofluorescence of yeast cells and growth media is substantially reduced with excitation wavelengths longer than 488 nm (Figure S1C).

To test whether these mCherry-greenFP fusions function as timers and to compare their ability to report on protein age, we analyzed cells expressing the proton pump Pma1 tagged with each putative tFT. Pma1 is an exceptionally stable protein of the plasma membrane (Thayer *et al.*, 2014). During yeast bud formation, preexisting Pma1 molecules at the plasma membrane are retained in the mother cell, while the bud (future daughter cell) receives newly produced Pma1 (Takizawa *et al.*, 2000; Malínská *et al.*, 2003). Consequently, cells expressing Pma1 tagged with the mCherry-sfGFP timer exhibit lower mCherry/greenFP intensity ratios (*R*) at the plasma membrane in the bud (*R*_b_) than in the mother cell (*R*_*m*_) (Figure 1C) (Khmelinskii *et al.*, 2012). We observed the same trend for all Pma1 fusions (Figure 1D), indicating that all tested mCherry-greenFP variants function as timers. The *R*_b_/*R*_m_ ratio can be used for a qualitative comparison of greenFP maturation rates with an approximate model of tFT maturation (Figure 1B) (Khmelinskii *et al.*, 2012). The *R*_b_/*R*_m_ ratio was lowest for mCherry-sfGFP and mCherry-mNeonGreen timers (Figure 1D), indicating that sfGFP and mNeonGreen have the fastest maturation among all tested greenFPs. Consistent with slower maturation of Clover (Figure S1A), the *R*_b_/*R*_m_ ratio was highest for the strain expressing Pma1-mCherry-Clover (Figure 1D). *De novo* fluorophore maturation of anaerobically synthesized GFPm is faster than maturation of sfGFP under the same conditions (Iizuka *et al.*, 2011) (Figure S1A). However, *in vivo* maturation (which also includes protein folding) of GFPm appears to be slower than that of sfGFP, as the *R*_b_/*R*_m_ ratio was higher for Pma1-mCherry-GFPm than for Pma1-mCherry-sfGFP (Figure 1D). The maturation kinetics of myeGFP is not known. The *R*_b_/*R*_m_ ratio of Pma1-mCherry-myeGFP was lower than expected for a greenFP with a maturation half-time of 25 min (i.e. the maturation half-time of EGFP (Figure S1A) (Shaner *et al.*, 2007)), indicating that myeGFP matures faster than EGFP but slower than sfGFP in yeast (Figure 1D).

We proceeded to examine the performance of all tFTs in comparative analysis of protein degradation kinetics. ln steady state, the mCherry/greenFP intensity ratio is determined by the degradation kinetics of the tFT fusion, such that strains expressing rapidly degraded fusions are expected to exhibit lower mCherry/greenFP intensity ratios (Figure 2A) (Khmelinskii *et al.*, 2012). We generated two constructs for expression of N-terminal ubiquitin (Ubi) fusions with each tFT (Ubi-X-mCherry-greenFP): one with a methionine residue (M) and another with an arginine residue (R) at position X (Figure 2B). Co-translational cleavage of the ubiquitin moiety exposes a new N-terminus starting with residue X, which determines the stability of the fusion according to the N-end rule. Fusions with N-terminal arginine, but not with N-terminal methionine, are recognized by the E3 ubiquitin ligase Ubr1 and targeted for proteasomal degradation (Bachmair *et al.*, 1986; Varshavsky, 2011) (Figure S2, A and B). Accordingly, for each tFT, the mCherry/greenFP intensity ratio was lower for the strain expressing Ubi-R-mCherry-greenFP (Figure 2C), indicating that all tested tFTs report on protein degradation kinetics.

**Figure 2.**
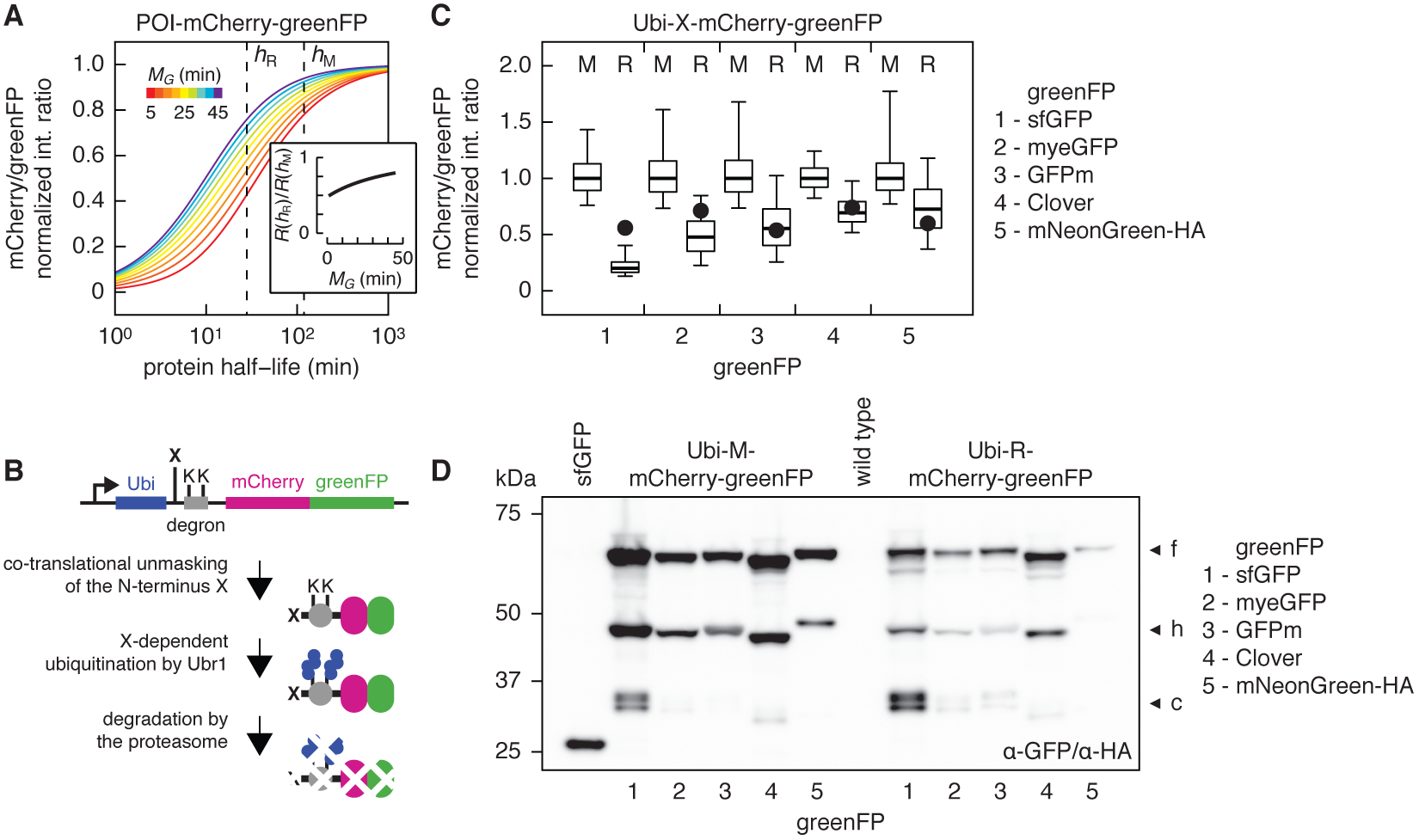
Analysis of protein degradation kinetics with different tFTs. A – Relationship between half-life of a tFT protein fusion and mCherry/greenFP intensity ratio in steady state. mCherry/greenFP intensity ratios were calculated as a function of protein degradation kinetics for a population doubling time of 90 min using published maturation parameters for mCherry (Khmelinskii *et al.*, 2012) and a maturation half-time *M*_G_ between 5 and 45 min for greenFP. Each curve is normalized to the mCherry/sfGFP intensity ratio of a non-degradable tFT fusion. Curves are color-coded according to *M*_G_ as indicated. Note that protein half-life *T* is related to degradation rate constant *k* as *T* = ln(2)/*k.* Similarly, maturation half-time *M*_G_ is related to maturation rate constant *m*_G_ as *M*_G_ = ln(2)/*m*_G_. Inset shows the comparison of mCherry/greenFP intensity ratios (*R*) for two protein half-lives – *h*_R_ = 28 min (half-life of R-mCherry-sfGFP) and *h*_M_ = 119 min (half-life of M-mCherry-sfGFP) (Figure S2B) – as a function of *M*_G_. B – Cartoon of Ubi-X-mCherry-greenFP constructs. Degradation rate of X-mCherry-greenFP fusions depends on the residue X, such that X = T (slow) < M < I < Y < F < R (fast). C – Fluorescence measurements with flow cytometry of strains expressing the indicated Ubi-X-mCherry-greenFP fusions. The residue X in each fusion is specified in the plot. Fusions with mNeonGreen carried a C-terminal HA tag to facilitate detection by immunoblotting. For each tFT, the mCherry/greenFP intensity ratios of individual cells were normalized to the median mCherry/greenFP intensity ratio of the corresponding Ubi-M-mCherry-greenFP fusion. Center lines mark the medians, box limits indicate the 25th and 75th percentiles, whiskers extend to 5th and 95th percentiles. Black circles mark theoretical mCherry/greenFP intensity ratios of Ubi-R-mCherry-greenFP (normalized to the corresponding Ubi-M-mCherry-greenFP fusions) calculated according to a simple model of mCherry-greenFP turnover in A using 28 and 119 min as half-lives of R-mCherry-sfGFP and M-mCherry-sfGFP, respectively, and published maturation half-times for mCherry and different greenFPs (Figure S1A). D – Immunoblot of strains expressing the indicated constructs. Whole cell extracts were separated by SDS-PAGE and probed with a mixture of antibodies against GFP and the HA tag. Three major forms observed for each Ubi-X-mCherry-greenFP fusion are indicated: a full-length X-mCherry-greenFP form (f), a shorter mCherry^ΔN^-greenFP product resulting from mCherry hydrolysis during cell extract preparation (h) (Gross *et al.*, 2000) and fast-migrating processed tFT fragments (c).

The difference between the median mCherry/greenFP intensity ratios of strains expressing Ubi-M-mCherry-greenFP and Ubi-R-mCherry-greenFP fusions varied from ~ 5 fold for the mCherry-sfGFP timer to 1.4 – 2.1 fold for the other tFTs. Comparison of these differences to an approximate model of tFT turnover (Figure 2A, Supplemental Theory) revealed several inconsistencies between theory and experiment. For instance, the relative mCherry/greenFP intensity ratio of Ubi-R-mCherry-sfGFP was considerably lower than expected (Figure 2C). Moreover, tFTs with Clover and mNeonGreen exhibited similar relative mCherry/greenFP intensity ratios for Ubi-R-mCherry-greenFP (Figure 2C), despite the difference in maturation kinetics (Figure S1A) and distinct performance as reporters of protein age (Figure 1D). These observations suggest that a parameter other than greenFP maturation kinetics influences tFT behavior in the protein degradation assay. Immunoblotting of whole cell extracts with antibodies against GFP (or against an hemagglutinin (HA) epitope fused to the C-terminus of mNeonGreen) revealed several bands in addition to full length fusions. First, a band at ~45 kDa (band (h) in Figure 2D) that is explained by hydrolysis of the mature mCherry fluorophore during cell extract preparation, which leads to a break in the mCherry polypeptide (Gross *et al.*, 2000; Shemiakina *et al.*, 2012). Additionally, unexpected tFT fragments were detected around 33 kDa (~6 kDa larger than free greenFP) in all strains expressing Ubi-X-mCherry-greenFP fusions (band (c) in Figure 2D). These fragments were most prominently seen for the tFT with sfGFP, especially for the Ubi-R-mCherry-sfGFP fusion. We sought to determine the origin of these protein fragments (hereafter referred to as processed tFT fragments) and to examine whether their presence can account for the unexpected behavior of different tFTs described above.

Two mechanisms could generate processed tFT fragments: proteolytic cleavage of the tFT by an endopeptidase (model 1) and incomplete degradation of the tFT by the proteasome (model 2) (Figure 3A). In model 1, the fraction of processed tFT in steady state 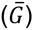 is expected to be independent of the degradation kinetics of the tFT fusion. In contrast, 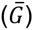 is expected to be proportional to the degradation kinetics of the tFT fusion in model 2 (further details in Supplemental Theory). In both models processed tFT fragments accumulate only if their degradation rate constant (*k*_2_ in Figure 3A) is small. Immunoblotting analysis of strains expressing a series of Ubi-X-mCherry-sfGFP fusions with different stabilities showed that the fraction of processed tFT increases as a function of protein degradation kinetics, in agreement with model 2 (Figure 3B). Moreover, accumulation of processed tFT fragments depended on proteasomal activity (Figure S2C). These observations were not specific to the Ubi-X-mCherry-greenFP fusions. Processed tFT fragments could also be detected in strains expressing misfolded cytoplasmic proteins tagged with mCherry-sfGFP and the fraction of processed tFT decreased upon deletion of E3 ubiquitin ligases involved in degradation of these misfolded proteins (Figure S2, D and E). Finally, similar ~33 kDa fragments have been observed when GFP fusions are degraded by proteasomes with impaired processivity, e.g. in mutants of the proteasome-associated E3 ubiquitin ligase Hul5 (or its mammalian ortholog UBE3C) (Zhang and Coffino, 2004; Aviram and Kornitzer, 2010; Martínez-Noël *et al.*, 2012; Chu *et al.*, 2013). We conclude that accumulation of processed tFT fragments is caused by incomplete proteasomal degradation of tFT fusions and that these fragments appear to be relatively stable in the cell.

**Figure 3.**
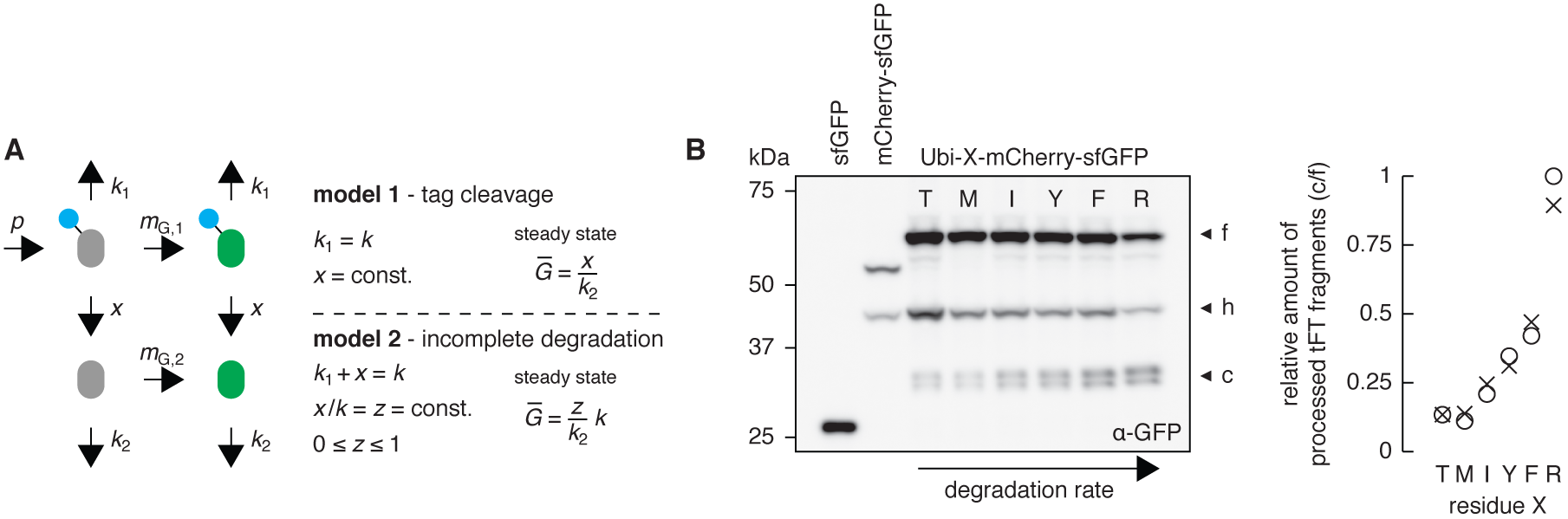
Incomplete degradation of tFT fusions leads to accumulation of processed tFT fragments. A – Turnover of a greenFP protein fusion with accumulation of processed greenFP. We assume that a greenFP protein fusion (protein of interest, represented by a blue circle, tagged with a greenFP) is produced at a constant rate *p* as a non-fluorescent protein and matures to a fluorescent protein in a single step with maturation rate constant *m*_G,1_. In model 1, greenFP protein fusions are degraded with rate constant *k* and greenFP can be cleaved off from both non-mature and mature protein fusions with rate constant *x*. In model 2, degradation of greenFP protein fusions with rate constant *k* proceeds to completion with probability 1 − z, such that greenFP fusions are effectively degraded with rate constant *k*_1_ = (1 – *z*)*k* and processed greenFP is produced with rate constant *x* = *kz*. Processed non-fluorescent greenFP matures in a single step with maturation rate constant *m*_G,2_ and is degraded with rate constant *k*_2_ in both models. In steady state, the fraction of processed greenFP (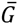 defined as the ratio between the total number of processed greenFP molecules and the total number of greenFP protein fusion molecules) is independent of *k* in model 1. However, 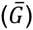 is proportional to *k* in model 2. Further details are provided as Supplemental Theory. B – Immunoblot of strains expressing the indicated constructs. The residue X in each Ubi-X-mCherry-sfGFP fusion is specified in the immunoblot. Three major forms observed for each Ubi-X-mCherry-sfGFP fusion are indicated as in Figure 2D. Note that in agreement with their degradation-dependent origin, processed tFT fragments were not detected in the strain expressing the stable mCherry-sfGFP fusion. The c/f ratio between the intensities of the (c) and (f) bands measured for each strain in two independent immunoblots, normalized to the maximum c/f ratio, is shown in the right panel.

What features of the mCherry-sfGFP timer are responsible for its incomplete degradation? We considered two possibilities: the linker connecting mCherry to sfGFP and the robust fold of sfGFP. The linker between mCherry and greenFP in the tFTs is composed of a leucine (L) and an aspartate (D) residues followed by five glycine-alanine (GA) repeats (LD(GA)_5_). GA rich sequences can prevent complete protein degradation by impairing the ability of the proteasome to unfold its substrate (Hoyt *et al.*, 2006; Daskalogianni *et al.*, 2008). To determine the role of the linker in the accumulation of processed tFT fragments, we tested shorter LDGAG or LDGS linker sequences between mCherry and sfGFP. Although shorter linkers changed the relative amounts of different processed tFT fragments, they did not reduce the total amount of processed fragments (Figure 4A) nor did they alter the ability of the mCherry-sfGFP timer to report on protein degradation kinetics (Figure 4B). In contrast, exchanging sfGFP for other greenFPs, while keeping the LD(GA)_5_ linker between mCherry and greenFP constant, reduced the amount of processed tFT fragments (Figure 2D). Moreover, fragments resulting from incomplete proteasomal degradation could be detected when using sfGFP but not mCherry alone as a tag (Figure S2F). This suggests that the stability of the greenFP fold, not the linker, is responsible for incomplete degradation of tFT fusions, although we cannot exclude the possibility that processed tFT fragments from timers other than mCherry-sfGFP are rapidly degraded in the cell.

**Figure 4.**
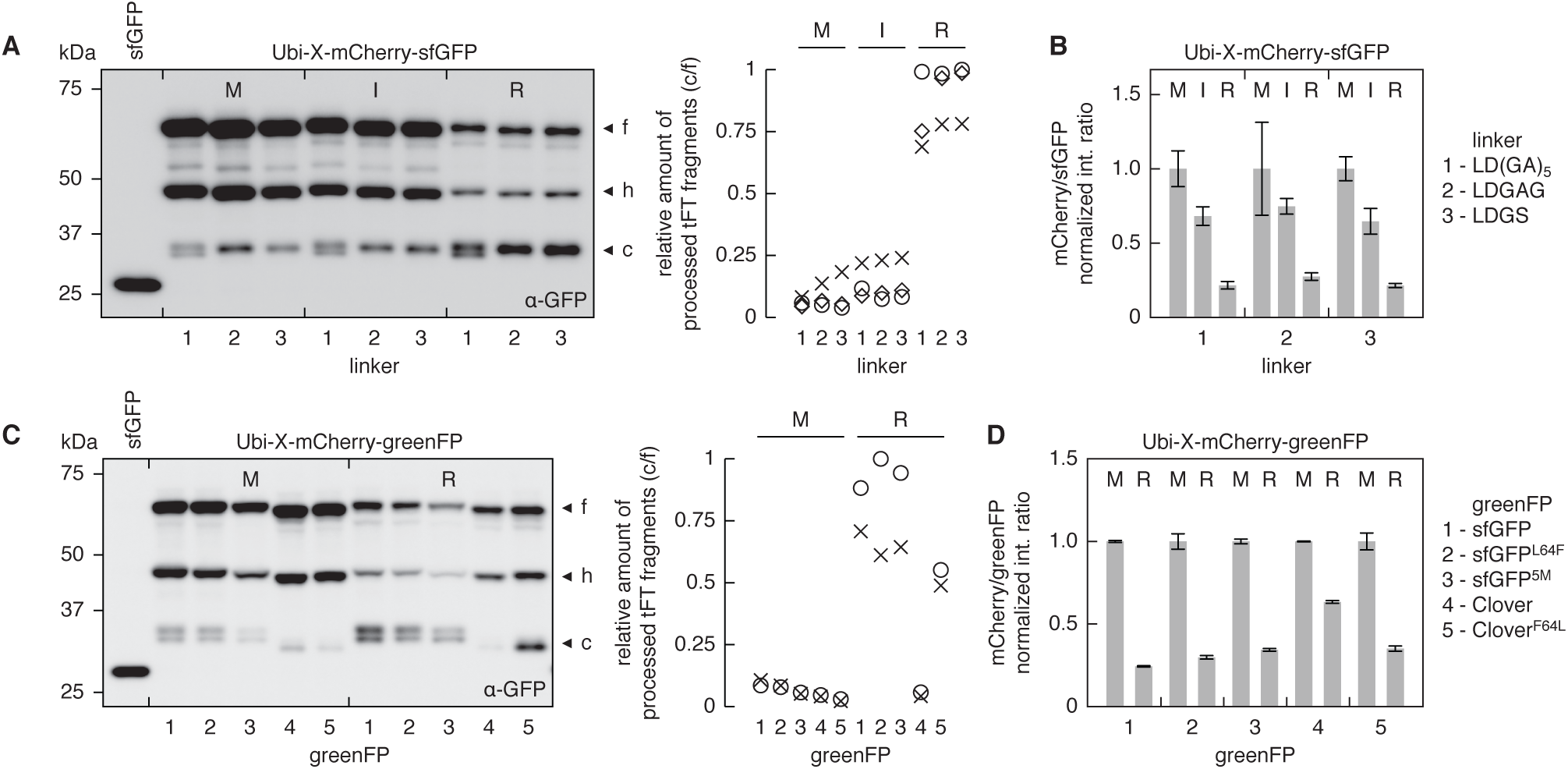
Stable greenFP fold prevents complete degradation of tFT fusions. A – Immunoblot of strains expressing Ubi-X-mCherry-sfGFP constructs with the indicated linkers between mCherry and sfGFP. The residue X in each Ubi-X-mCherry-sfGFP fusion is specified in the immunoblot. Whole cell extracts were separated by SDS-PAGE and probed with antibodies against GFP. Three major forms observed for each Ubi-X-mCherry-sfGFP fusion are indicated as in Figure 2D. The c/f ratio between the intensities of the (c) and (f) bands measured for each strain in three independent immunoblots, normalized to the maximum c/f ratio, is shown in the right panel. B – mCherry/sfGFP intensity ratios determined from whole colony fluorescence measurements of strains expressing Ubi-X-mCherry-sfGFP constructs in A. For each linker, mCherry/sfGFP intensity ratios (mean ± s.d., *n* ≥ 12 per construct) were normalized to the corresponding Ubi-M-mCherry-sfGFP fusion. C – Immunoblot of strains expressing Ubi-X-mCherry-greenFP constructs with the indicated greenFPs. The residue X in each Ubi-X-mCherry-greenFP fusion is specified in the immunoblot. Three major forms observed for each fusion are indicated as in Figure 2D. The c/f ratio between the intensities of the (c) and (f) bands measured for each strain in two independent immunoblots, normalized to the maximum c/f ratio, is shown in the right panel. D – mCherry/sfGFP intensity ratios determined from whole colony fluorescence measurements of strains expressing Ubi-X-mCherry-greenFP constructs in C. For each greenFP, mCherry/greenFP intensity ratios (mean ± s.d., *n* ≥ 4 per construct) were normalized to the corresponding Ubi-M-mCherry-greenFP fusion.

To test the possibility that incomplete degradation is caused by the robust fold of sfGFP, we attempted to destabilize the sfGFP fold by reverting five mutations (S30R, Y39N, N105T, Y145F and I171V) responsible for the superfolder nature of sfGFP (Pédelacq *et al.*, 2006) (sfGFP^5M^) or by reverting the F64L mutation known to improve GFP folding at 37°C (Cormack *et al.*, 1996; Tsien, 1998) (sfGFP^L64F^). However, tFTs with these sfGFP mutants behaved similarly to the mCherry-sfGFP timer (Figure 4, C and D). Nevertheless, in a reverse experiment, introducing the F64L mutation into Clover increased the accumulation of processed tFT fragments in the strain expressing Ubi-R-mCherry-Clover^F64L^ (Figure 4C). This accumulation correlated with a reduced mCherry/greenFP intensity ratio of Ubi-R-mCherry-Clover^F64L^ relative to the Ubi-M-mCherry-Clover^F64L^ fusion, bringing it close to Ubi-R-mCherry-sfGFP (Figure 4D). The F64L mutation did not affect the accumulation of processed tFT fragments for the Ubi-M-mCherry-Clover^F64L^ fusion, possibly because the longer half-life of Ubi-M-tFT fusions gives enough time for complete folding and maturation of wild type Clover. These observations support the idea that the stability of the greenFP fold is responsible incomplete degradation of tFT fusions.

Next we sought to destabilize the sfGFP fold through circular permutations. Circular permutation can drastically impair folding of classical GFP (Baird *et al.*, 1999; Topell *et al.*, 1999). Although sfGFP is largely tolerant of circular permutations (Pédelacq *et al.*, 2006), some circularly permutated sfGFP variants can be efficiently unfolded and more rapidly degraded by the AAA+ ClpXP protease *in vitro* (Nager *et al.*, 2011). We constructed a series of tFTs with different circular permutations (cp) of sfGFP (Figure 5A) and examined their behavior in the degradation assay with N-terminal ubiquitin fusions. Accumulation of processed tFT fragments was reduced to below the detection limit with all circular permutations except cp3 (Figure 5B). This is consistent with higher mobility of the β_7_-β_11_ sheets in the β-barrel fold and the tendency of GFP to start unfolding from β_7_-β_11_ (Huang *et al.*, 2007; Zimmer *et al.*, 2014). As seen in the case of Clover (Figure 4, C and D), reduced accumulation of processed tFT fragments correlated with a reduced relative difference in mCherry/sfGFP ratios between the stable and unstable Ubi-X-mCherry-greenFP fusions (Figure 5C). Taken together, these experiments indicate that the stability of the greenFP fold is responsible for proteasome-dependent processing of tFTs and concomitant accumulation of tFT fragments.

**Figure 5.**
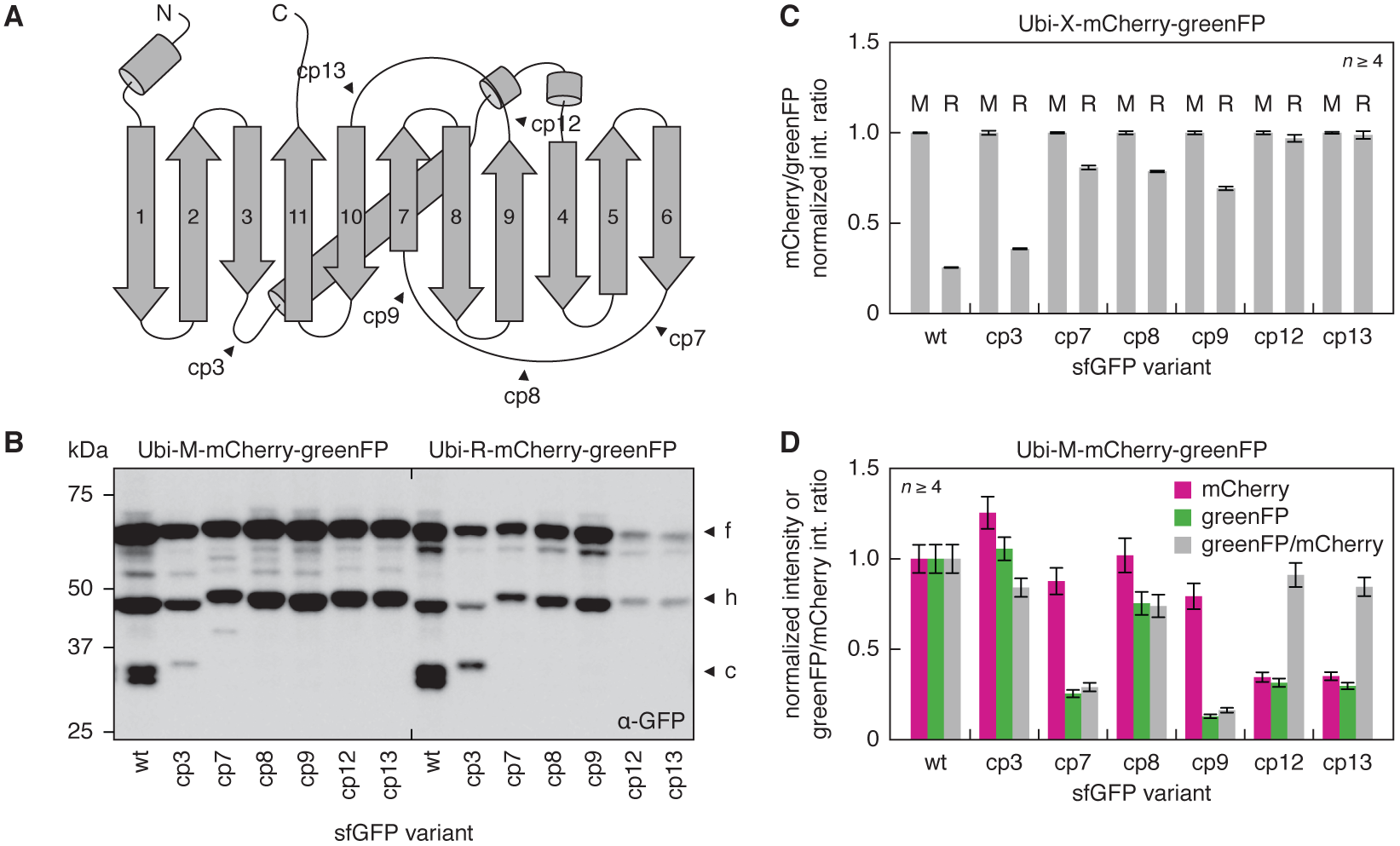
Improved degradation of tFT fusions with circular permutations of sfGFP. A – Schematic representation of sfGFP. Start sites of six circular permutations (cp) are indicated, numbered according to (Pédelacq *et al.*, 2006). B – Immunoblot of strains expressing Ubi-X-mCherry-greenFP constructs with different circular permutations of sfGFP as greenFP. Three major forms observed for each fusion are indicated as in Figure 2D. C, D – Whole colony fluorescence measurements of strains expressing Ubi-X-mCherry-greenFP constructs in A. C – The residue X in each fusion is specified in the plot. For each sfGFP variant, mCherry/greenFP intensity ratios (mean ± s.d., *n* ≥ 4 per construct) were normalized to the corresponding Ubi-M-mCherry-greenFP fusion. D – mCherry fluorescence intensities (measure of expression levels), greenFP fluorescence intensities and greenFP/mCherry intensity ratios (measure of greenFP molecular brightness) (mean ± s.d., *n* ≥ 4 per construct) were normalized to Ubi-M-mCherry-sfGFP.

Two permutations, cp7 and cp9, reduced the molecular brightness of sfGFP (Figure 5D). Another two, cp12 and cp13, appeared to affect protein folding and expression levels (Figure 5D), and the maturation kinetics of sfGFP, as suggested by the lack of difference in mCherry/sfGFP ratios between the two Ubi-X-mCherry-greenFP fusions (Figure 5C). However, the cp8 permutation did not impair protein expression and had only a minor effect on the molecular brightness of sfGFP (Figure 5D). sfGFP(cp8) exhibited the fastest maturation kinetics among all tested circular permutation, as evidenced by the performance of the mCherry-sfGFP(cp8) timer in the analysis of Pma1 protein age (Figure 1D). Moreover, accumulation of processed tFT fragments was strongly reduced in human embryonic kidney HEK293T cells when using the mCherry-sfGFP(cp8) timer instead of mCherry-sfGFP (Figure S3). Thus, sfGFP(cp8) represents an alternative to sfGFP without potential artifacts caused by incomplete proteasomal degradation.

We sought to determine how incomplete proteasomal degradation affects the properties of tFTs and their use in studies of protein dynamics. We incorporated proteasome-dependent processing into a model of tFT maturation and turnover for two tFTs, mCherry-sfGFP and sfGFP-mCherry. Proteasomal degradation typically requires an unstructured region in the substrate to initiate degradation (Prakash *et al.*, 2004). We assumed that no such initiation region is present in the mCherry-sfGFP and sfGFP-mCherry timers. It is implicit in the model that degradation is initiated within the tagged protein of interest and proceeds in a processive manner until the sfGFP fold is reached, at which point the remaining polypeptide can be either released from the proteasome with a defined probability or completely degraded. Incomplete degradation of proteins tagged at the C-terminus with mCherry-sfGFP or sfGFP-mCherry produces respectively free sfGFP or free sfGFP-mCherry in the model (further details in Supplemental Theory). Our experiments indicate that the mCherry-sfGFP timer functions as a degradation reporter with C-terminally tagged proteins (Figure 2) (Khmelinskii *et al.*, 2012). However, the time range of this timer shifts towards more stable proteins when the probability of incomplete tFT degradation is increased in the model (Figures 6A). This shift does not depend on whether Förster resonance energy transfer (FRET) from sfGFP to mCherry is considered in the model (Supplemental Theory and Figure S4). The difference in mCherry/sfGFP intensity ratios between Ubi-M-mCherry-sfGFP and Ubi-R-mCherry-sfGFP fusions is expected to increase with increasing probability of incomplete tFT degradation (Figure 6A). The experimentally observed behavior of Clover and sfGFP-based tFTs with different levels of incomplete tFT degradation is consistent with this prediction (Figures 4D and 5C).

**Figure 6.**
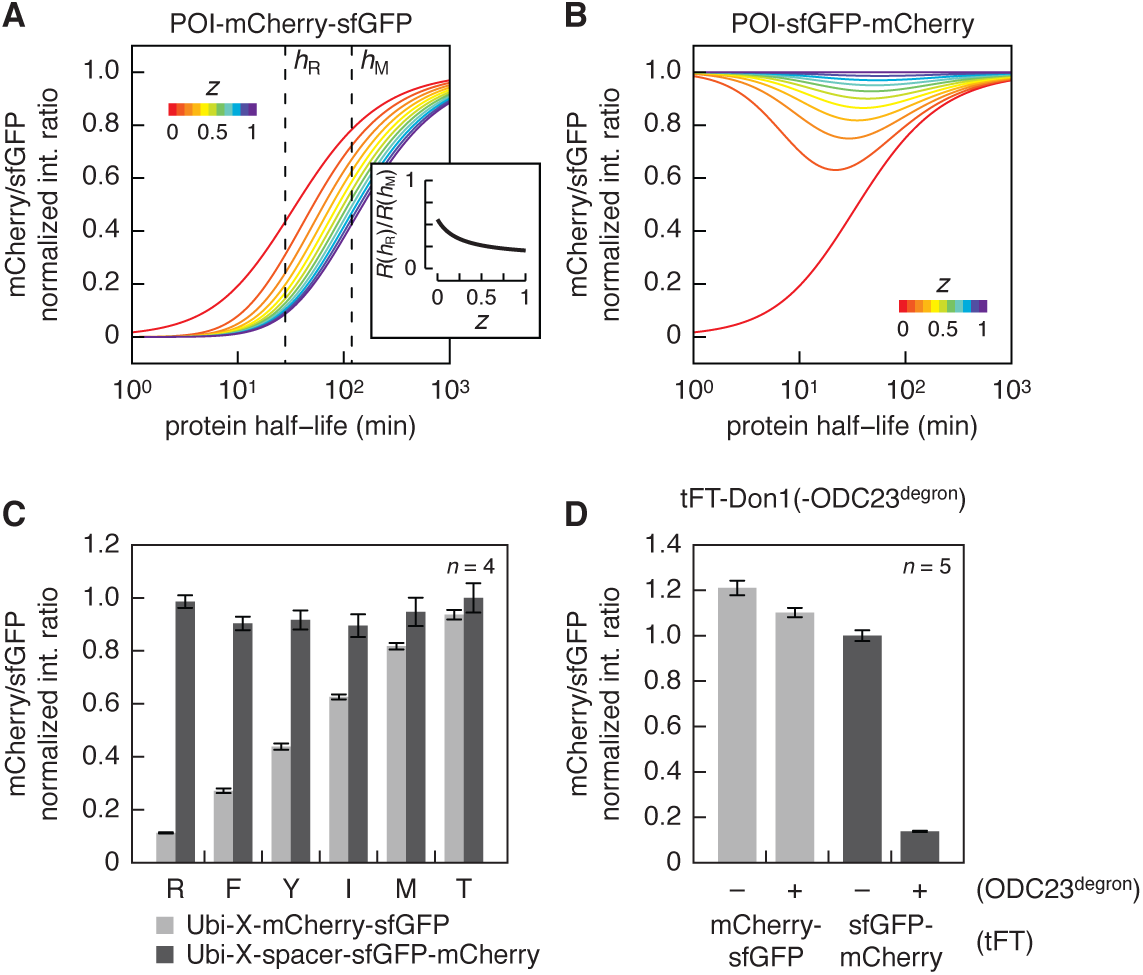
Proteasome-dependent processing constrains the design of tFTs. A, B – Relationship between half-life of a protein of interest (POI) tagged at the C-terminus with a tFT and mCherry/sfGFP intensity ratio in steady state. mCherry/sfGFP intensity ratios were calculated as a function of protein degradation kinetics for a population doubling time of 90 min using published maturation parameters for mCherry and sfGFP (Khmelinskii *et al.*, 2012) and a probability *z* of incomplete degradation of sfGFP between 0 and 1. Incomplete degradation produces free sfGFP in A or free sfGFP-mCherry in B. Each curve is normalized to the mCherry/sfGFP intensity ratio of a non-degradable tFT fusion. Curves are color-coded according to *z* as indicated. Further details are provided as Supplemental Theory. Inset shows the comparison of mCherry/sfGFP intensity ratios (*R*) for two protein half-lives – *h*_R_ = 28 min (half-life of R-mCherry-sfGFP) and *h*_M_ = 119 min (half-life of M-mCherry-sfGFP) (Figure S2B) – as a function of *z*. C – Fluorescence measurements with flow cytometry of strains expressing the indicated Ubi-X-tFT fusions. The residue X in each fusion is specified in the plot. mCherry/sfGFP intensity ratios (mean ± s.d., *n* = 4 per construct) were normalized to Ubi-T-spacer-sfGFP-mCherry. D – Whole colony fluorescence measurements of strains expressing tFT-Don1 fusions with and without an ODC23 degron at the C-terminus. The tFT in each fusion is specified in the plot. mCherry/sfGFP intensity ratios (mean ± s.d., *n* = 5 per construct) were normalized to sfGFP-mCherry-Donl.

In contrast to the mCherry-sfGFP timer, incomplete degradation of the sfGFP-mCherry timer as a C-terminal tag is predicted to abolish the monotonic relationship between mCherry/sfGFP intensity ratio and protein half-life (Figure 6B). We tested this prediction in the degradation assay with N-terminal ubiquitin fusions. To ensure initiation of degradation N-terminally to sfGFP, we separated the N-terminal degradation signal (N-degron) from the sfGFP-mCherry timer with a spacer protein Don1 and an HA tag (Ubi-X-spacer-sfGFP-mCherry). Immunoblotting of whole cell extracts confirmed the accumulation of processed tFT fragments with the expected size, ~6 kDa larger than free sfGFP-mCherry, in these strains (Figure S5A). In contrast to strains expressing Ubi-X-mCherry-sfGFP fusions, the mCherry/sfGFP intensity ratio was largely independent of the degradation rate for Ubi-X-spacer-sfGFP-mCherry fusions (Figures 6C and S5B). This result is consistent with the model predictions and suggests that the probability of incomplete sfGFP degradation is substantial (~0.5 or higher) (Figure 6B). Moreover, it shows that the sfGFP-mCherry timer should not be used as a degradation reporter with C-terminally tagged proteins. However, it is important to stress that the sfGFP-mCherry fusion is a timer, i.e. it changes color with time, and thus should report on the age of C-terminally tagged proteins in the absence of protein degradation. Indeed cells expressing Pma1-sfGFP-mCherry exhibited lower mCherry/sfGFP intensity ratios at the plasma membrane in the bud than in the mother cell, similarly to Pma1-mCherry-sfGFP (Figure 1D). Moreover, the sfGFP-mCherry timer, but not the mCherry-sfGFP timer, faithfully reported on the degradation kinetics of N-terminally tagged proteins (Figures 6D and S5C). Together, these results indicate that different tFT variants should be used for N- and C-terminal protein tagging, such that the sfGFP moiety is placed distally to the region in the protein of interest where proteasomal degradation is initiated. Importantly, this theoretical analysis was performed under the assumption that processed tFT fragments are stable in the cell (Figure 6, A and B), and thus establishes an upper limit for the influence of proteasome-dependent processing of tFTs on their ability to report on protein degradation kinetics. Increasing the degradation rate of processed tFT fragments reduces their impact and, at the limit of infinitely fast degradation, mCherry-sfGFP and sfGFP-mCherry timers are both reporters of protein degradation kinetics when used for C-terminal tagging, with behavior indistinguishable from a hypothetical mCherry-sfGFP timer that is completely degraded by the proteasome (Supplemental Theory and Figure S6).

## Discussion

We report here on the use of different greenFPs for analysis of protein dynamics with tFTs. All tested greenFPs can be combined with mCherry to obtain tFTs that report on protein age and degradation kinetics, indicating that greenFPs mature faster than mCherry. Our analysis shows that the greenFP maturation kinetics influences the time range of mCherry-greenFP timers (Figure 1). In addition, we observe that greenFPs (especially sfGFP), but not mCherry, can withstand proteasomal degradation in yeast (Figures 2 and S2F), consistent with previous reports of incomplete proteasomal degradation of GFP (Zhang and Coffino, 2004; Aviram and Kornitzer, 2010; Martínez-Noël *et al.*, 2012; Chu *et al.*, 2013). The efficiency of FP degradation by the proteasome could differ in other organisms and cell types, as proteasome processivity varies between species (Kraut *et al.*, 2012).

Two features are necessary for a protein to become a proteasome substrate: a degradation signal (degron) and a degradation initiation site (Schrader *et al.*, 2009). These features are provided by the model substrates used to investigate tFT behavior in the degradation assay (Figure 2). Therefore, proteasomal degradation of a tFT starts from the site where it is linked to the model substrate and proceeds by unfolding and channeling of the polypeptide into the catalytic core of the 19S particle of the proteasome. Protein sequences that cannot be efficiently channeled into the catalytic core can be release from the proteasome. Several factors can contribute to this release: the protein sequence adjacent to the released domain, the stability of the fold of the released domain, and an intrinsic probability of the proteasome to release substrates that is independent of the power stroke underlying substrate unfolding (Kraut *et al.*, 2012; Fishbain *et al.*, 2015). Our analysis indicates that incomplete degradation of sfGFP fusions depends on the stability of the sfGFP fold (Figures 4 and 5), although the molecular details, i.e. the determinants in sfGFP, are unclear.

The outcome of proteasomal processing of tFT fusions depends on the order of FPs in the timer relative to the region in the protein from which proteasomal degradation begins. If mCherry is placed proximal to the degradation initiation site (e.g. in mCherry-sfGFP timer as a C-terminal tag), proteasomal processing produces tFT fragments of ~33 kDa (Figure 2). These contain the greenFP moiety and a short ~6 kDa tail that presumably corresponds to the length of the unfolded polypeptide channeled into the proteasome catalytic core before release of the fragments. In this situation, the timer functions as a reporter of protein age and degradation kinetics but its time range is shifted towards more stable proteins compared to a hypothetical timer that is completely degraded by the proteasome (Figure 6A). This shift should increase the difference in mCherry/greenFP intensity ratios between the stable Ubi-M-tFT and unstable Ubi-R-tFT fusions (inset in Figure 6A), as experimentally observed for the mCherry-sfGFP timer compared to mCherry-sfGFP(cp8) or mCherry-mNeonGreen (Figures 2C and 5C). Moreover, the increased cytoplasmic greenFP fluorescence resulting from incomplete tFT degradation should be taken into account when measuring protein age at subcellular level. We note that the ~33 kDa tFT fragments are distinct from the ~26 kDa fragments observed upon incomplete lysosomal/vacuolar degradation of GFP fusions, which have been exploited to follow the cytoplasm to vacuole targeting (Cvt) pathway and autophagy (Shintani and Klionsky, 2004; Klionsky *et al.*, 2008). If in turn greenFP is located proximal to the degradation initiation site, proteasomal processing results in the release of ~60 kDa tFT fragments containing both FPs and a short ~6 kDa tail (Figure S5A). In this case the timer can be still used to measure local protein age but it no longer reports on protein degradation kinetics (Figure 6B). From these observations we can derive a short list of recommendations for design and use of tFTs as reporters of protein age and degradation:

i. The time range of a tFT is mostly determined by the maturation kinetics of the slower-maturing FP (Khmelinskii *et al.*, 2012), although the maturation kinetics of the faster-maturing FP, the extent of proteasomal processing and potential FRET between the two FPs also contribute (Figures 2A, 6A and S4A). Thus, to construct a tFT for a given time range, a slower-maturing FP with maturation kinetics matching the time scale of interest should be paired with a faster-maturing FP that is as bright and fast-maturing as possible to lower the detection limit of the timer (Khmelinskii and Knop, 2014).
ii. Ideally, characterization of a new tFT should involve testing both arrangements of FPs as reporters of protein age and protein degradation in the experimental system of interest to obtain qualitative system-specific estimates of FP maturation rates, efficiency of proteasomal degradation and time range of the tFT. In yeast, this can be done using the Pma1 protein age and Ubi-X-tFT degradation assays (Figures 1D and 2C).
iii. For tFTs based on mCherry and greenFPs (or any timer in which one of the FPs is resistant to proteasomal degradation), the FPs should be arranged such that the degradation-resistant greenFP ends up distal to the degradation initiation site in the tagged protein of interest. Thus, the mCherry-sfGFP timer should be used for C-terminal tagging, whereas the sfGFP-mCherry timer – for N-terminal tagging. For internal tagging, both timers should be tested to first identify the degradation initiation site based on the size of processed tFT fragments.
iv. Generally, a tFT in which both FPs are efficiently degraded by the proteasome is preferable, assuming its time range is suitable for the intended analysis. However, tFTs with incomplete proteasomal degradation can be advantageous in some situations. For instance, although the time range of mCherry-sfGFP(cp8) and mCherry-mNeonGreen timers is in principle more suitable to study degradation of unstable proteins than mCherry-sfGFP (Figure 6A), the mCherry-sfGFP timer could facilitate detection of extremely unstable and low abundance proteins due to increased absolute greenFP fluorescence that comes from accumulation of processed tFT fragments. Moreover, processed tFT fragments can be exploited as a marker of proteasomal degradation. For example, for a protein tagged at the C-terminus with mCherry-sfGFP, accumulation of ~33 kDa fragments is indicative of proteasomal degradation (Figure S2C), whereas lysosomal/vacuolar degradation should produce ~26 kDa fragments (Shintani and Klionsky, 2004; Klionsky *et al.*, 2008).
v. tFTs with incomplete proteasomal degradation such as mCherry-sfGFP can be used for systematic analysis of protein degradation as the mCherry/sfGFP intensity ratio in steady state is a monotonic function of protein half-life (Figure 6A). However, care should be exercised when comparing proteins degraded by the proteasome and lysosome/vacuole pathways, which produce distinct tFT degradation fragments and differ in local environment that can affect FP maturation and brightness (Khmelinskii and Knop, 2014).

To follow protein dynamics, fast-maturing FP tags are needed to detect proteins as early as possible after synthesis. Our analysis indicates that mNeonGreen (Lam *et al.*, 2012) is a good alternative to sfGFP: it is similarly fast-maturing but significantly less resistant to proteasomal degradation in yeast than sfGFP (Figures 1 and 2). The molecular brightness of mNeonGreen is similar to sfGFP when using excitation/emission wavelengths optimal for sfGFP and ~5 times higher with wavelengths optimal for mNeonGreen (Figure S1). The only limitation is that filter sets matching the excitation/emission peaks of mNeonGreen or lasers around its 506 nm excitation peak are not commonly used in fluorescence microscopy. The cp8 circular permutation of sfGFP is another alternative. When used as a C-terminal tag, sfGFP(cp8) is completely degraded by the proteasome both in yeast and human cells, and its maturation kinetics and brightness are similar to sfGFP (Figures 5 and S3).

In conclusion, our study provides a detailed characterization of tFTs as reporters of protein age and degradation. This should facilitate their application in different areas of cellular and organismal research. Our work also emphasizes the notion that fluorescent proteins are not neutral tags. Careful characterization of their properties is required for correct interpretation of protein dynamics measurements across spatial and temporal dimensions.

## Materials and Methods

### Yeast methods and plasmids

Yeast genome manipulations (gene deletions and tagging) were performed using conventional PCR targeting, as described (Janke *et al.*, 2004). Yeast strains and plasmids used in this study are listed in Supplemental Tables 1 and 2, respectively. Yeast codon-optimized sequences of all fluorescent proteins were obtained using full gene synthesis. All Ubi-X-tFT constructs are based on previously described Ubi-X-β-galactosidase fusions, where X is followed by a 40-residue sequence that starts with histidine (Bachmair *et al.*, 1986). The sequence of the ubiquitin-independent ODC23 degron (MSCAQESITSLYKKAGSENLYFQ) was obtained from plasmid pCT334 (Renicke *et al.*, 2013). The Don1 coding sequence was amplified from genomic DNA of strain ESM356-1 (Supplemental Table 1). Standard site-directed mutagenesis was used to introduce point mutations into sfGFP or Clover and change the linker between mCherry and sfGFP. Circular permutations of sfGFP were amplified from a plasmid carrying two copies of sfGFP fused with a short linker (GSGAG). The N-termini of circular permutations 3, 7, 8, 9, 12 and 13 are the amino acids at position 51, 129, 140, 145, 189 and 189 of sfGFP, respectively. All plasmid sequences are available upon request.

### Models of tFT maturation and turnover

Fluorescence intensity curves depicting maturation of a pool of mCherry-sfGFP molecules initialized in the non-mature state in the absence of protein production and degradation (Figure 1A) were calculated using a two-step maturation model for mCherry (maturation half-times of 16.91 and 30.3 min) and one-step maturation model for sfGFP (maturation half-time of 5.63 min), as described (Khmelinskii *et al.*, 2012). To examine the influence of greenFP on tFT maturation (Figure 1B), the maturation half-time of greenFP was varied between 5 and 45 min in 5 min steps. All curves were normalized to the point of complete maturation.

To examine the influence of greenFP maturation on the relationship between protein half-life and mCherry/greenFP intensity ratio in steady state (Figure 2A), mCherry/greenFP intensity ratios were calculated according to equation E33 (Supplemental Theory), using a probability of incomplete tFT degradation *z* = 0, mCherry maturation half-times of 16.91 and 30.3 min, greenFP maturation half-time between 5 and 45 min varied in 5 min steps and a population doubling time of 90 min.

To examine the influence of incomplete tFT degradation on the relationship between protein half-life and mCherry/sfGFP intensity ratio in steady state (Figure 6, A and B), mCherry/sfGFP intensity ratios were calculated for the mCherry-sfGFP and sfGFP-mCherry timers according to equations E33 and E34, respectively (Supplemental Theory), using a population doubling time of 90 min, mCherry maturation halftimes of 16.91 and 30.3 min for both full-length fusion and processed form, sfGFP maturation half-time of 5.63 min for both full-length fusion and processed form, a degradation rate constant of processed tFT fragments *k*_2_ = 0 and varying the probability of incomplete tFT degradation *z* between 0 and 1 in steps of 0.1.

To examine the influence of FRET within a tFT on the relationship between protein half-life and mCherry/sfGFP intensity ratio in steady state (Figure S4A), mCherry/sfGFP intensity ratios were calculated for the mCherry-sfGFP timer according to equation E39 (Supplemental Theory), using a population doubling time of 90 min, a probability of incomplete tFT degradation *z* = 0, mCherry maturation half-times of 16.91 and 30.3 min, sfGFP maturation half-time of 5.63 min and varying the FRET efficiency *E* between 0 and 1 in steps of 0.1. For a fixed FRET efficiency *E* (0, 0.5 or 1) (Figures S4, B-D), the calculations were extended by varying the probability of incomplete tFT degradation *z* between 0 and 1 in steps of 0.1, using a degradation rate constant of processed tFT fragments *k*_2_ = 0 and sfGFP maturation half-time of 5.63 min for both full-length fusion and processed form.

To examine how degradation of processed tFT fragments influences the relationship between protein half-life and mCherry/sfGFP intensity ratio in steady state (Figure S6), mCherry/greenFP intensity ratios were calculated for the mCherry-sfGFP and sfGFP-mCherry timers according to equations E33 and E34, respectively (Supplemental Theory), using a population doubling time of 90 min, mCherry maturation halftimes of 16.91 and 30.3 min for both full-length fusion and processed form, sfGFP maturation half-time of 5.63 min for both full-length fusion and processed form, a probability of incomplete tFT degradation *z* = 1 and varying the half-life of processed tFT fragments *H*_2_ between 10 and 1000 min, in addition to 0 and infinity. The half-life of processed tFT fragments *H*_2_ is related to the degradation rate constant of processed tFT fragments *k*_2_ as *k*_2_ = ln(2)/*H*_2_.

### Fluorescence microscopy

Strains were grown at 30°C in low fluorescence medium (synthetic complete medium prepared with yeast nitrogen base lacking folic acid and riboflavin (CYN6501, ForMedium)) to 0.4−1.2×10^7^ cells ml−^1^ and attached to glass-bottom 96-well plates (MGB096-1-2-LG-L, Matrical) using Concanavalin A (C7275, Sigma) as described (Khmelinskii and Knop, 2014). Images from the middle of the cell were acquired on a DeltaVision Elite system (Applied Precision), consisting of an inverted epifluorescence microscope (IX71; Olympus) equipped with an LED light engine (SpectraX, Lumencor), 475/28 and 575/25 excitation, and 525/50 and 624/40 emission filters (Semrock), a dual-band beam splitter 89021 (Chroma Technology), a 100x NA 1.4 UPlanSApo oil immersion objective (Olympus), an sCMOS camera (pco.edge 4.2, PCO) and a motorized stage contained in a temperature-controlled chamber. Image correction and quantification were performed using ImageJ (Schneider *et al.*, 2012). Dark signal and flat field corrections were applied to all images as described (Khmelinskii and Knop, 2014). Outlines of mother and bud compartments were manually defined in the sfGFP channel using a 5 pixel wide segmented line with a spline fit and applied to the mCherry channel. Fluorescence measurements at the plasma membrane were corrected for background using autofluorescence of the growth medium measured in close proximity to each individual cell.

### Flow cytometry

Strains were grown at 30°C in synthetic medium lacking leucine (to select for plasmids) to a density of 0. 4−1.2×10^7^ cells ml^−1^. Single cell fluorescence intensities, forward and side scatter were measured for at least 6000 cells per sample on a BD FACSCanto RUO Special Order System (BD Biosciences) equipped with 488 nm and 561 nm lasers, 505 nm and 600 nm long pass filters, 530/30 nm and 610/20 nm band pass filters. Multi-spectral beads (3.0-3.4 μm Sphero Rainbow Calibration Particles (6 peaks), #556286, BD Biosciences) were used to control for fluctuations in excitation laser power. Data analysis was performed with Flowing Software 2 www.flowingsoftware.com): measurements were gated for cells, followed by gating for cells with fluorescence above background to exclude cells that lost the expression plasmids. Sample measurements were corrected for background using autofluorescence levels of a control strain carrying an empty plasmid. mCherry/sfGFP intensity ratios were calculated for each individual cell.

### Colony fluorescence measurements

Strains were grown to saturation at 30°C in synthetic medium lacking leucine and pinned onto agar plates with synthetic medium lacking leucine in 384-colony format, with 4 technical replicates for each strain. For pinning a RoToR pinning robot (Singer Instruments) was used. Fluorescence intensities of colonies were typically measured after 24 h of growth at 30°C using an Infinite M1000 Pro plate reader (Tecan). Measurements in mCherry (587/10 nm excitation, 610/10 nm emission, optimal detector gain) and sfGFP (488/10 nm excitation, 510/10 nm emission, optimal detector gain) channels were performed from the top at 400 Hz frequency of the flash lamp, with 20 flashes averaged for each measurement. For relative measurements of molecular brightness of different greenFPs (Figure S1B), fluorescence intensities were additionally measured in a third channel (505/5 nm excitation, 516/5 nm emission, optimal detector gain). Measurements of control colonies without fluorescent protein expression were used to correct all measurements for background.

To acquire excitation spectra (Figure S1C), exponentially growing cultures of strains yMaM26 and yMaM32 (Supplemental Table 1) were transferred to glass-bottom 96-well plates (MGB096-1-2-LG-L, Matrical) and allowed to settle down. Fluorescence emission at 510/5 nm was measured from the bottom with excitation wavelengths between 350 and 600 nm, in 2 nm steps, at 400 Hz frequency of the flash lamp, with 1 flash per step.

### Immunoblotting and time-course experiments

Strains were typically grown at 30°C in synthetic medium lacking leucine to 10^7^ cells ml^−1^. One-milliliter samples were mixed with 150 *μ*l of 1.85 M NaOH and 10 *μ*l β-mercaptoethanol, and flash frozen in liquid nitrogen. Whole cell extracts were prepared as previously described (Knop *et al.*, 1999), separated by SDSPAGE (NuPAGE Novex 4-12% Bis-Tris protein gels (Life Technologies)) followed by blotting and probed with mouse anti-HA (12CA5), mouse anti-myc (9E10), rabbit anti-GFP (ab6556, abcam), rabbit anti-YFP (Miller *et al.*, 2015), mouse anti-Pgk1 (22C5D8, Molecular Probes) and rabbit anti-Zwf1 (Miller *et al.*, 2015) antibodies. Secondary goat anti-mouse (IgG (H+L)-HRP, Dianova) and goat anti-rabbit (IgG (H+L)-HRP, Dianova) antibodies were used for detection. Imaging was performed on a LAS-4000 system (Fuji) using the increment exposure routine to avoid overexposure. Image quantification was performed in ImageJ (Schneider *et al.*, 2012). All measurements were corrected for local background.

For cycloheximide chases, strains were grown at 30°C in synthetic complete medium to ~0.8×10^7^ cells ml^−1^ before addition of cycloheximide to 100 *μ*g/ml final concentration. One-milliliter samples taken at each time point were immediately mixed with 150 *μ*l of 1.85 M NaOH and 10 *μ*l β-mercaptoethanol, flash frozen in liquid nitrogen and processed for immunoblotting as detailed above.

For the proteasome inhibition experiment (Figure S2C), strains yMaM67 and yMaM957 (Supplemental Table 1) were grown at 30°C to ~0.8×10^7^ cells ml^−1^ in synthetic medium lacking leucine and with raffinose (2% w/v) as carbon source. Expression of Ubi-R-mCherry-sfGFP was induced by addition of galactose (2% w/v) alongside inhibition of proteasome activity by addition of MG132 (40 mM stock in DMSO) to 80 *μ*g/ml final concentration or DMSO as control. Whole cell extracts of samples collected before and after induction were prepared and analyzed by SDS-PAGE followed by immunoblotting, as detailed above.

### Human cell culture and immunoblotting

The sequence encoding PARK6 was amplified from the plasmid pcDNA3.1/Pink1 (Meissner *et al.*, 2011) and cloned into the pcDNA5/mCherry-sfGFP vector (Khmelinskii *et al.*, 2012). pcDNA5/PARK6-mCherry-sfGFP(cp8) was cloned by overlap assembly PCR of a human codon-optimized sfGFP(cp8), which was obtained by full gene synthesis, into the pcDNA5/PARK6-mCherry-sfGFP background.

HEK293T cells were grown in Dulbecco’s modified Eagle’s medium supplemented with 10% (v/v) fetal bovine serum at 37°C in 5% (v/v) CO2. Transient transfections were performed using 25 kDa linear polyethylenimine (Polysciences) (Durocher *et al.*, 2002) and harvested 24 h thereafter. To block mitochondrial protein import, cells were incubated for 3 h with 10 *μ*M carbonyl cyanide *m*-chlorophenyl hydrazone (Sigma) before cell lysis.

Whole cell extracts were separated by SDS-PAGE followed by blotting onto a PVDF membrane and probed with a monoclonal anti-GFP antibody (11814460001, Roche). A secondary goat anti-mouse IgG-HRP antibody (sc-2005, Santa Cruz) was used for detection. Imaging was performed on a LAS-4000 system (Fuji).

### Supplemental Materials

Supplemental Materials contain Supplemental Theory, Supplemental Figures S1-S6 and Supplemental Tables 1 and 2.

## Acknowledgements

We thank Christof Taxis for reagents and Marc Zimmer for comments on the manuscript. This work was supported by the Deutsche Forschungsgemeinschaft (DFG) through the Sonderforschungsbereich 1036 (SFB1036) (M.K., A.M., B.B. and M.K.L.) and the Hartmut Hoffmann-Berling International Graduate School of Molecular and Cellular Biology (HBIGS) (C.H.).Figure 1. Analysis of protein age with different tFTs.

## Supplemental Materials

### 1. Supplemental Theory

#### 1.1. Turnover of sfGFP fusions with tag cleavage

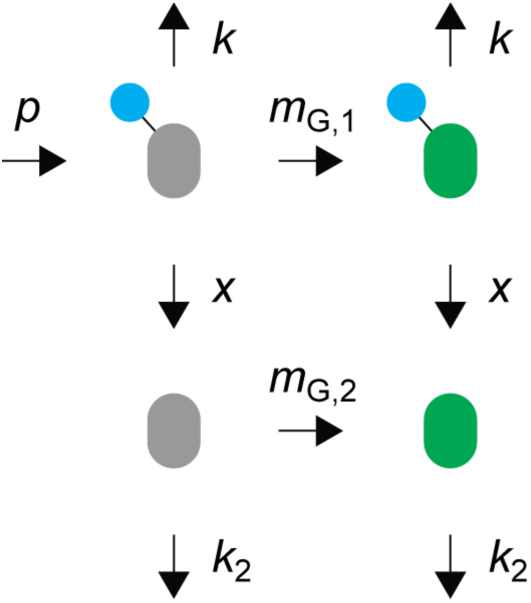

In this model (model 1 in Figure 3A) we assume that a fluorescent protein fusion, a protein of interest (blue in the cartoon) tagged with sfGFP (grey or green), is produced at a constant rate *p* as a non-fluorescent protein and matures to a fluorescent protein in a single step with the maturation rate constant *m*_G,1_. Both non-mature and mature protein fusions are degraded with the rate constant *k.* Additionally, sfGFP can be cleaved off from both non-mature and mature protein fusions with the rate constant *x*. Non-fluorescent cleaved sfGFP matures in a single step with the maturation rate constant *m*_G,2_ and both forms of cleaved sfGFP are degraded with the rate constant *k*_2_.

As a result of one-step maturation and tag cleavage, there are four populations of protein species: a population of non-mature (non-fluorescent) protein fusions with *N_d_* members, a population of mature (fluorescent) protein fusions with *N_d_* members, a population of non-mature cleaved sfGFP with *N_m_* members and a population of mature cleaved sfGFP with 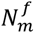 members. The following rate equations describe the dynamics of these four populations:

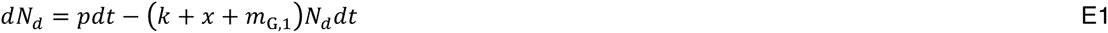

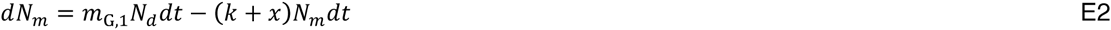

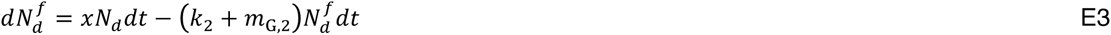

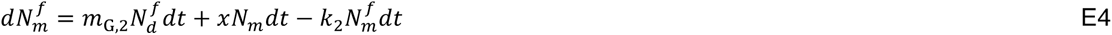

The steady-state solution of the set of differential equations E1-E4 is as follows:

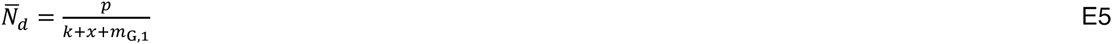

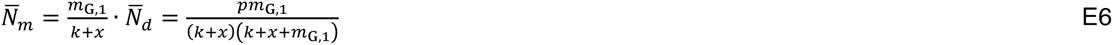

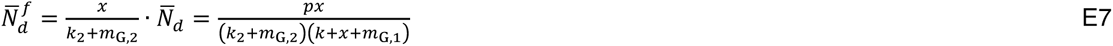

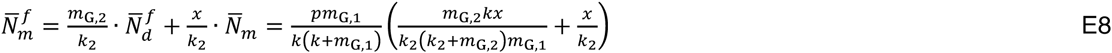

The ratio 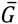 between the number of cleaved sfGFP molecules and full-length fusions in steady state is therefore given by:

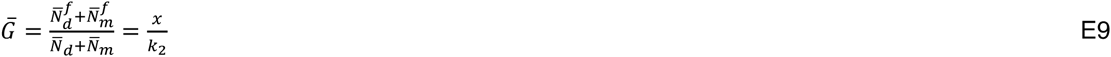

#### 1.2. Turnover of sfGFP fusions with incomplete degradation of sfGFP

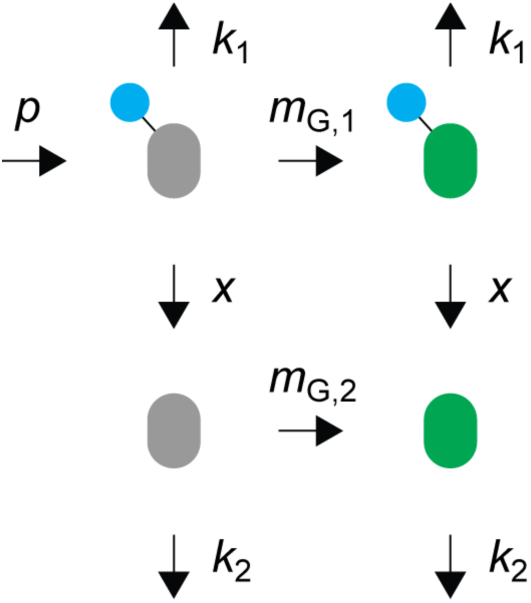

In this model (model 2 in Figure 3A) we assume that a fluorescent protein fusion (protein of interest tagged with sfGFP) is produced at a constant rate *p* as a non-fluorescent protein and matures to a fluorescent protein in a single step with the maturation rate constant *m*_G,1_. Both non-mature and mature protein fusions are degraded with the rate constant *k* but degradation proceeds to completion with probability (1 – *z*), such that full-length fusions are effectively degraded with the rate constant = (1 – z)*k* and, as a result, processed (free) sfGFP is produced with the rate constant *x* = *kz*. Non-fluorescent processed sfGFP matures in a single step with the maturation rate constant *m*_G,2_ and both forms of processed sfGFP are degraded with the rate constant *k*_2_. We assume that the probability of incomplete degradation *z* is a property of the sfGFP tag (i.e. it does not depend on the protein of interest) and by definition has a value between 0 and 1.

As a result of one-step maturation and incomplete degradation, there are four populations of protein species: a population of non-mature (non-fluorescent) protein fusions with *N_d_* members, a population of mature (fluorescent) protein fusions with *N_m_* members, a population of non-mature processed sfGFP with members and a population of mature processed sfGFP with 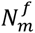 members. The following rate equations describe the dynamics of these four populations:

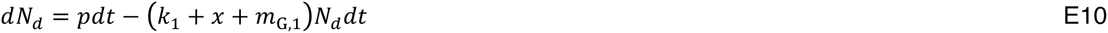

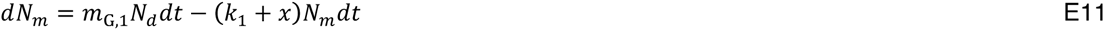

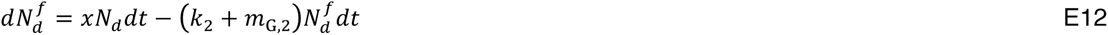

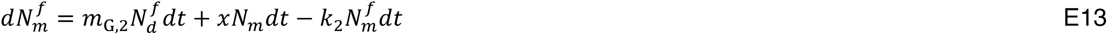

The steady-state solution of the set of differential equations E10-E13 is as follows:

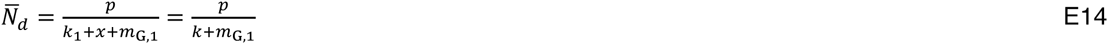

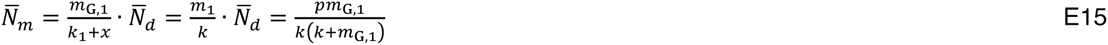

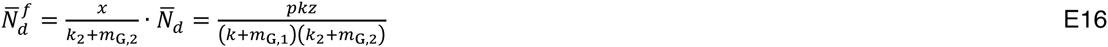

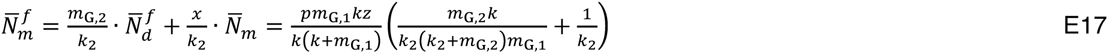

Note that 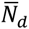 and 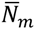 populations are independent of incomplete degradation, in agreement with our assumptions above.

The total number 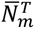 of fluorescent molecules in steady state is therefore given by:

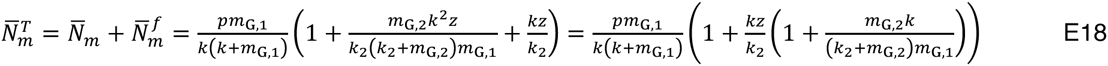

And the steady-state ratio 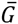 between the number of processed sfGFP molecules and full-length fusions is given by:

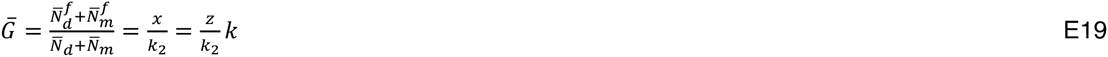

#### 1.3. Turnover of mCherry fusions with incomplete degradation of mCherry

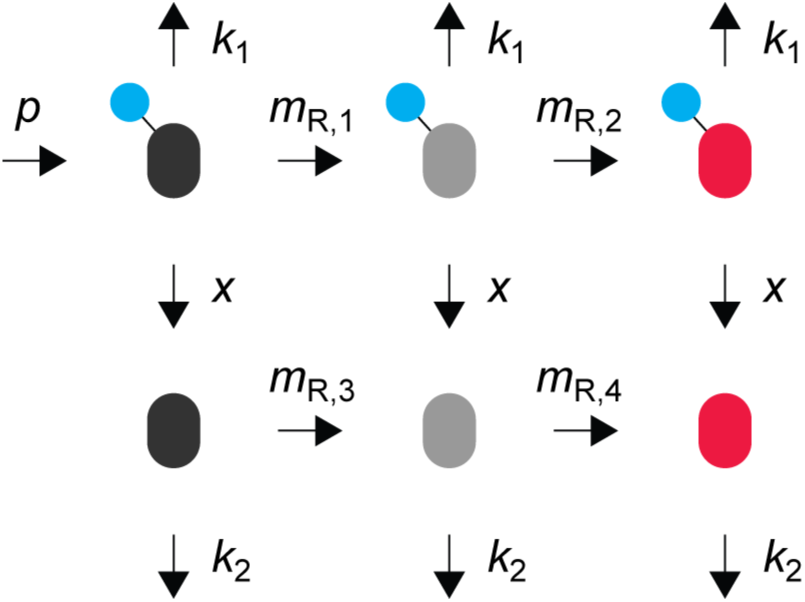

In this model we assume that a fluorescent protein fusion (protein of interest tagged with mCherry) is produced at a constant rate *p* as a non-fluorescent protein and matures to a fluorescent protein in a two-step process with the maturation rate constants *m*_R,1_ and *m*_R,2_. Both non-mature and mature protein fusions are degraded with the rate constant *k* but degradation proceeds to completion with probability (1 – *z*), such that full-length fusions are effectively degraded with the rate constant *k*_1_ = (1 – *z*)*k* and processed mCherry is produced with the rate constant *x* = *kz.* Non-fluorescent processed mCherry matures in a two-step process with the maturation rate constants *m*_R,3_ and *m*_R,4_. All forms of processed mCherry are degraded with the rate constant *k*_2_. We assume that the probability of incomplete degradation *z* is a property of the mCherry tag (i.e. it does not depend on the protein of interest) and by definition has a value between 0 and 1.

As a result of two-step maturation and incomplete degradation, there are six populations of protein species: a population of non-mature (non-fluorescent) protein fusions with *N_d_* members (dark grey in the cartoon), a population of protein fusions in the intermediate maturation state with *N_i_* members (light grey), a population of mature (fluorescent) protein fusions with *N_m_* members, a population of non-mature processed mCherry with 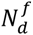 members (dark grey), a population of processed mCherry in the intermediate maturation state with 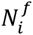 members (light grey) and a population of mature processed mCherry with 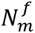 members. The following rate equations describe the dynamics of these six populations:

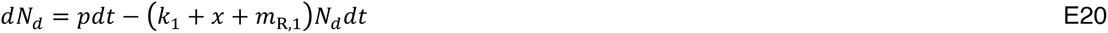

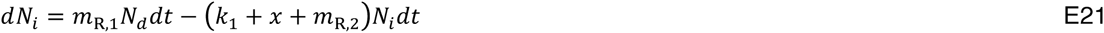

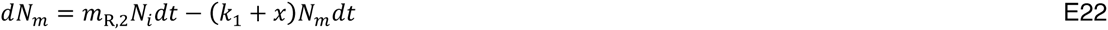

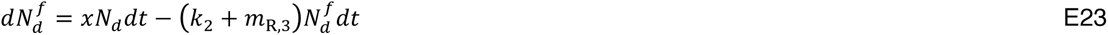

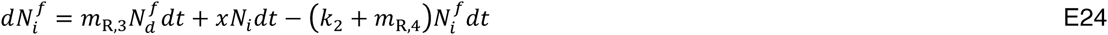

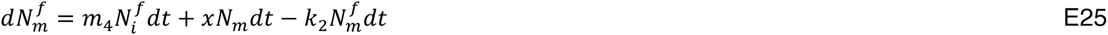

The steady-state solution of the set of differential equations E20-E25 is as follows:

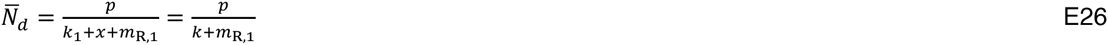

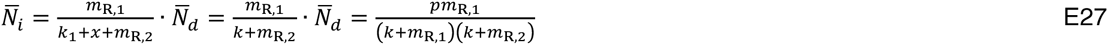

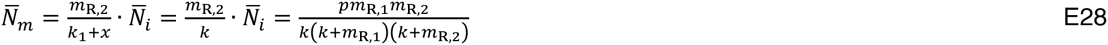

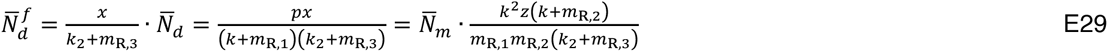

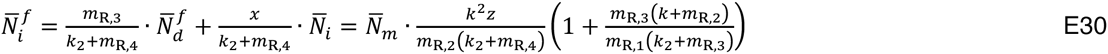

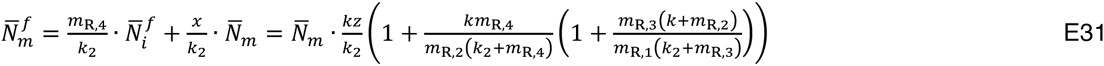

Note that 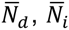 and 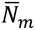 populations are independent of incomplete degradation, in agreement with our assumptions above.

The total number 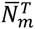 of fluorescent molecules in steady state is therefore given by:

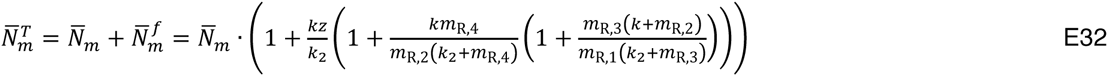

#### 1.4. Turnover of mCherry-sfGFP fusions with incomplete degradation of sfGFP

Here we consider the maturation and turnover of a protein tagged at the C-terminus with the mCherry-sfGFP timer (or tagged at the N-terminus with the sfGFP-mCherry timer). The fusion is produced at a constant rate *p* as a non-fluorescent protein. The two fluorescent proteins in the tag mature independently: mCherry undergoes a two-step maturation with rate constants *m*_R,1_ and *m*_R,2_ whereas sfGFP matures in a single step with the maturation rate constant *m*_G,1_. We assume that degradation is initiated within the tagged protein of interest and proceeds in a processive manner until the sfGFP fold is reached, at which point the remaining polypeptide can be released from the proteasome with probability *z* or completely degraded with probability (1 − *z*). Therefore, full-length fusions are effectively degraded with the rate constant *k_i_* = (1 − *z*)*k* and processed (free) sfGFP is produced with the rate constant *x = kz*. Non-fluorescent processed sfGFP matures in a single step with the maturation rate constant *m*_G,1_ and both forms of processed sfGFP are degraded with the rate constant *k*_2_. We assume that the probability of incomplete degradation *z* is a property of the sfGFP in the tag (i.e. it does not depend on the protein of interest) and by definition has a value between 0 and 1.

In steady state, the population of fluorescent sfGFP molecules 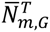 is given by equation E18, whereas the population of fluorescent mCherry molecules 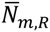 is given by equation E28. The steady-state ratio Γ_1.4_ of fluorescent populations 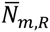 and 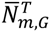 is therefore given by:

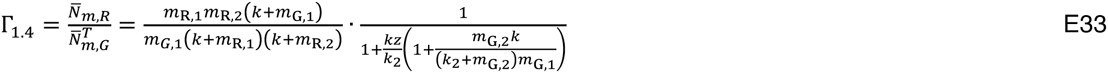

The ratio Γ_1.4_ is independent of the rate of protein production, as previously observed for tFT turnover without incomplete degradation of sfGFP (Khmelinskii *et al.*, 2012). Assuming that *k*_2_, the degradation rate constant of processed sfGFP, is independent of the protein of interest tagged with the tFT (i.e. incomplete proteasomal degradation of tFT fusions always leads to production of the same processed sfGFP fragments), the ratio Γ_1.4_ depends on only one variable, the degradation rate constant *k* of the tagged protein of interest. Note that for z = 0, Γ_1.4_ is reduced to the steady-state ratio 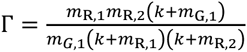 derived for the case of tFT turnover without incomplete degradation of tFT fusions (Khmelinskii *et al.*, 2012). Similarly, Γ_1.4_ = Γ when degradation of processed sfGFP is extremely fast (*k*_2_ *→* ∞).

#### 1.5. Turnover of sfGFP-mCherry fusions with incomplete degradation of sfGFP-mCherry

Here we consider the maturation and turnover of a protein tagged at the C-terminus with the sfGFP-mCherry timer (this model also applies to proteins tagged at the N-terminus with the mCherry-sfGFP timer). The fusion is produced at a constant rate *p* as a non-fluorescent protein. The two fluorescent proteins in the tag mature independently: mCherry undergoes a two-step maturation with rate constants *m*_R,1_ and *m*_R,2_ whereas sfGFP matures in a single step with the maturation rate constant *m*_G,1_. We assume that degradation is initiated within the tagged protein of interest and proceeds in a processive manner until the sfGFP fold is reached, at which point the remaining polypeptide can be released from the proteasome with probability z or completely degraded with probability (1 − z). Therefore, full-length fusions are effectively degraded with the rate constant *k_i_* = (1 − z)*k* and processed sfGFP-mCherry (free tFT) is produced with the rate constant *x* = *kz.* The two fluorescent proteins in the free tFT mature independently – mCherry undergoes a two-step maturation with rate constants *m*_R,3_ and *m*_R,4_ whereas sfGFP matures in a single step with the maturation rate constant *m*_G,2_ – and all forms of processed sfGFP-mCherry are degraded with the rate constant *k*_2_. We assume that the probability of incomplete degradation *z* is a property of the sfGFP in the tag (i.e. it does not depend on the protein of interest) and by definition has a value between 0 and 1.

In steady state, the population of fluorescent sfGFP molecules 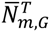 is given by equation E18, whereas the population of fluorescent mCherry molecules 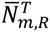 is given by equation E32. The steady-state ratio Γ_1.5_ of fluorescent populations 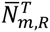 and 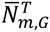 is therefore given by:

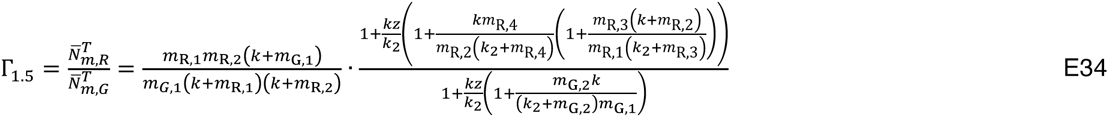

The ratio Γ_1.5_ is independent of the rate of protein production. Assuming that *k*_2_ is independent of the protein of interest tagged with the tFT, the ratio Γ_1.5_ depends on only one variable, the degradation rate constant *k* of the tagged protein of interest. Note that Γ_1.4_ = Γ_1.5_ = Γ when degradation of tFT fusions always proceeds to completion (z = 0) or when degradation of processed tFT is extremely fast (*k*_2_ → ∞).

#### 1.6. Turnover of mCherry-sfGFP fusions with incomplete degradation of sfGFP, with FRET

Here we extend the model describing turnover of mCherry-sfGFP fusions with incomplete degradation of sfGFP (section 1.4) to include the possibility of Förster resonance energy transfer (FRET) between the two fluorophores in the tFT. We assume that FRET is possible from sfGFP to mCherry within the tFT (but not from free sfGFP to mCherry) such that excitation of an sfGFP fluorophore can results in fluorescence emission by an mCherry fluorophore instead of the initially excited sfGFP fluorophore. This energy transfer can occur because the excitation spectrum of the acceptor mCherry fluorophore overlaps with the emission spectrum of the donor sfGFP fluorophore and the two are sufficiently close in space. The probability by which the energy absorbed by the sfGFP fluorophore is transferred to the mCherry fluorophore is the FRET efficiency *E.*

In steady state, the fluorescence intensity *I_R_* of mCherry using an excitation wavelength that only allows direct excitation of mCherry fluorophores is FRET-independent and proportional to the number 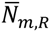 of mature mCherry fluorophores given by equation E28:

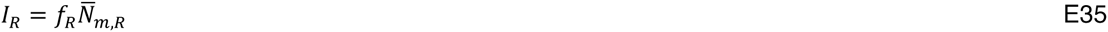

In contrast, the fluorescence intensity *Ĩ_G_* of sfGFP is FRET-dependent and has contributions from three distinct states of sfGFP. The first contribution is not affected by FRET and comes from free sfGFP molecules. The second contribution is also not affected by FRET and comes from tFT molecules with mature sfGFP but non-mature mCherry. The third contribution comes from tFT molecules with mature sfGFP and mature mCherry fluorophores. In this case the sfGFP fluorescence is reduced due to FRET as a fraction *E* of the total excitation energy is transferred from sfGFP to mCherry. By defining *b* as the steady-state probability for a neighbor of a mature sfGFP fluorophore being a mature mCherry fluorophore, *Ĩ_G_* can be expressed as follows: 

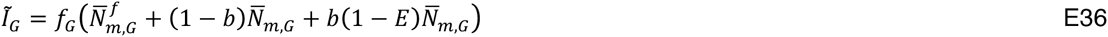
where 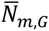 and 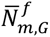 are given by equations E15 and E17, respectively. By substituting the instrument-specific proportionality constant *f = f_R_/f_G_*, the FRET-depedent intensity ratio 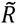 results as: 

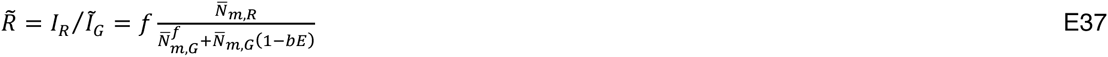
where the probability *b* is defined as:

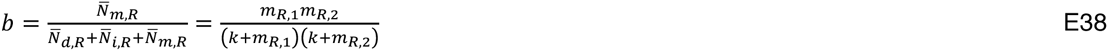

By substituting *b,* we obtain the FRET-depedent intensity ratio 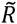:

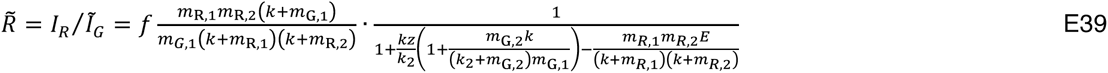

In the absence of FRET, i.e. *E =* 0, the intensity ratio 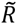 is reduced to the FRET-independent ratio *R = f*Γ_1.4_.

### 2. Supplemental Figures

#### 2.1. Supplemental Figure S1

**Supplemental Figure S1.**
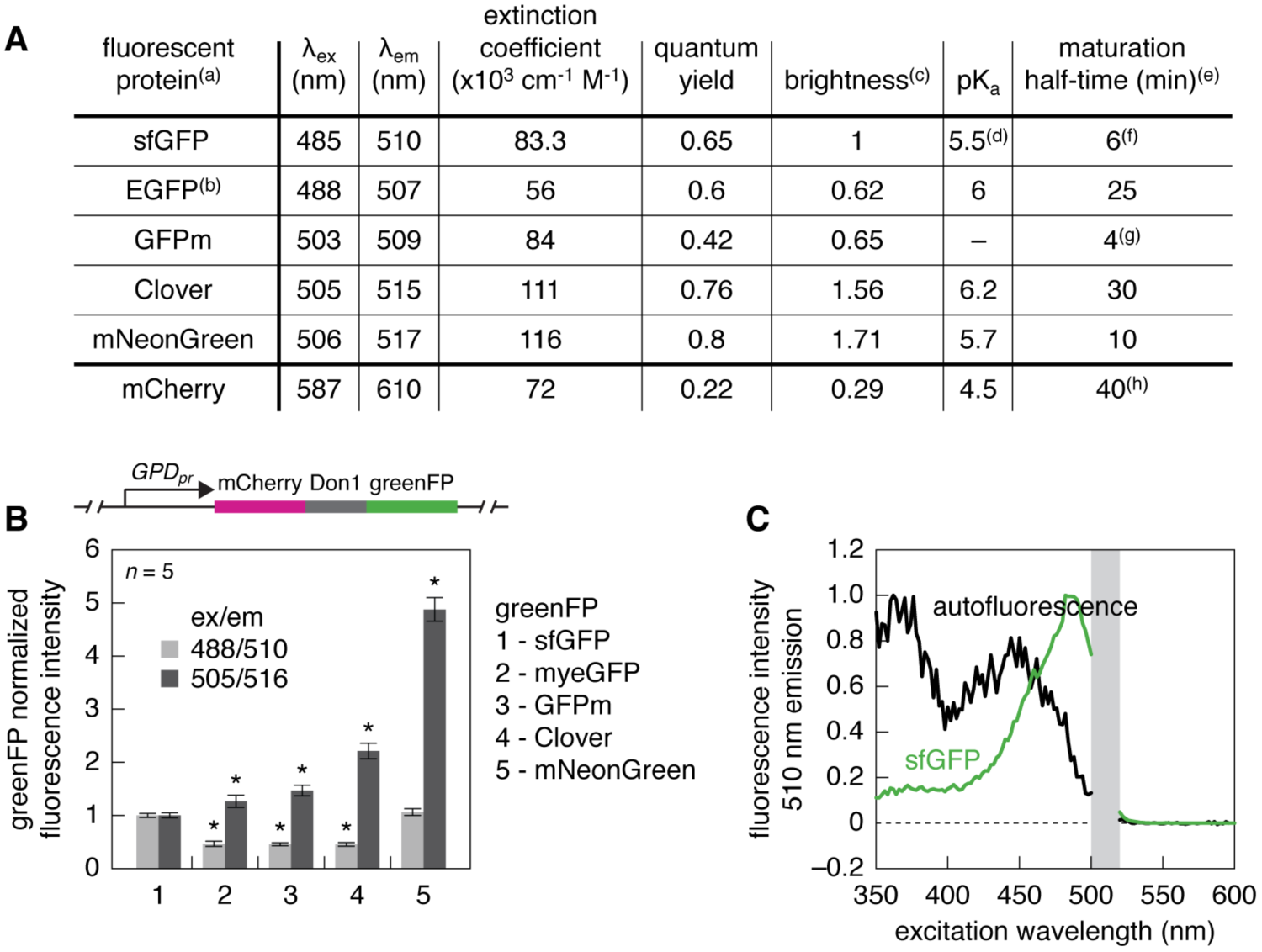
Properties of different greenFPs. A – Properties of fluorescent proteins used in this study. Notes: (a) – values were obtained from the following references unless indicated otherwise: sfGFP (Pédelacq *et al.*, 2006), EGFP (Shaner *et al.*, 2007), GFPm (Yoo *et al.*, 2007; Iizuka *et al.*, 2011), Clover (Lam *et al.*, 2012), mNeonGreen (Shaner *et al.*, 2013), mCherry (Shaner *et al.*, 2004); (b) – properties of myeGFP have not been measured, EGFP is shown instead; (c) – brightness, defined as the product of extinction coefficient and quantum yield, is normalized to sfGFP; (d) – value obtained from (Shaner *et al.*, 2013); (e) – the methods used for maturation half-time measurements vary between publications, therefore the values cannot be compared to each other; (f) – maturation of sfGFP was measured *in vivo* in yeast (Khmelinskii *et al.*, 2012); (g) – maturation of GFPm was measured upon exposure of anaerobically produced protein to oxygen, for comparison sfGFP and EGFP maturation halftimes measured using this assay are ~16 and ~9.6 min, respectively (Iizuka *et al.*, 2011); (h) – maturation of mCherry *in vivo* in yeast follows a two-step process with half-times of 17 and 30 min (Khmelinskii *et al.*, 2012). B – Measurements of molecular brightness of different greenFPs in yeast. Each greenFP was fused to mCherry with Don1 as a spacer protein to minimize Förster resonance energy transfer (FRET) between the two fluorescent proteins. The fusions were expressed in yeast from the strong constitutive *GPD* promoter. Whole colony greenFP fluorescence intensities, measured with two sets of excitation (ex) and emission (em) wavelengths, were normalized for protein expression levels using mCherry fluorescence intensities. The resulting estimates of molecular brightness (mean ± s.d., *n* = 5 biological replicates each with 4 technical replicates) were normalized to sfGFP. Significant differences from sfGFP molecular brightness are indicated (^*^, p < 0.002 in a two-tailed t-test). C – Excitation spectra of yeast strains with and without sfGFP expression. Each spectrum is normalized to the maximum fluorescence intensity. Note the substantial levels of colony autofluorescence upon excitation at 488 nm.

#### 2.2. Supplemental Figure S2

**Figure S2.**
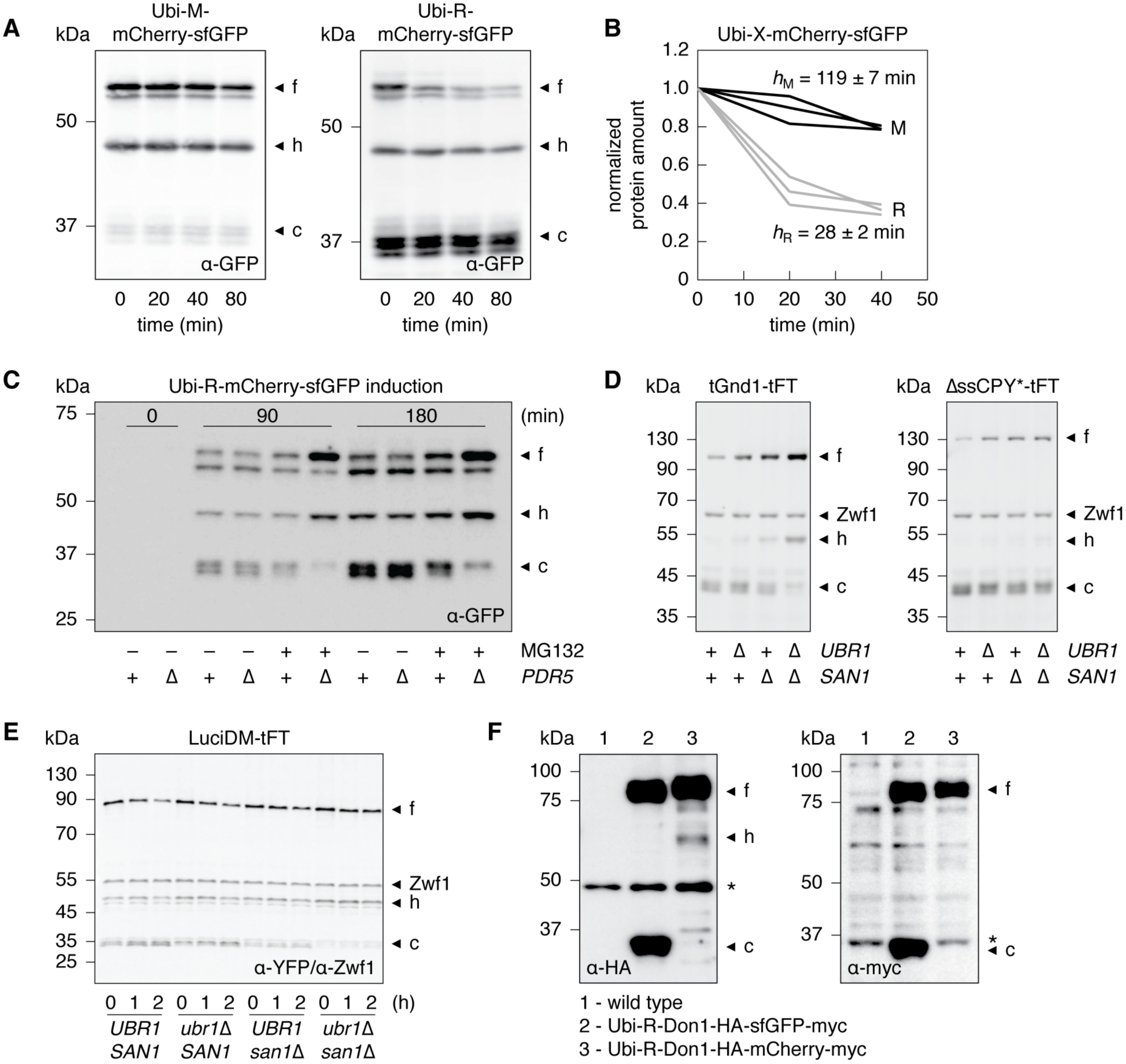
Accumulation of processed tFT fragments results from incomplete proteasomal degradation of tFT fusions. A – Degradation of Ubi-X-mCherry-sfGFP fusions after blocking translation with cycloheximide. Whole cell extracts of samples collected at the indicated time points were separated by SDS-PAGE and probed with antibodies against GFP. Three major forms observed for each Ubi-X-mCherry-sfGFP fusion are indicated as in Figure 2D. B – Quantification of protein half-lifes for the Ubi-X-mCherry-sfGFP fusions. The residue X in each Ubi-X-mCherry-sfGFP fusion is specified in the plot. The amount of the (f) form was measured in 3 replicates of the cycloheximide chase experiment in A and normalized to the starting amount for each replicate. The estimated half-lives (*h*_x_, mean ± s.d.) are indicated. C – Expression of Ubi-R-mCherry-sfGFP was induced in a wild type strain (wt) or in a strain sensitized to chemical inhibition of proteasome activity through deletion of *PDR5* (Δ), a gene encoding a multidrug transporter. The proteasome inhibitor MG132 or dimethyl sulfoxide (DMSO, as control) were added concomitantly with the start of induction. Whole cell extracts of samples collected at the indicated time points after induction were separated by SDS-PAGE. Three major forms detected with antibodies against GFP are indicated as in Figure 2D. D – Immunoblots of strains expressing truncated Gnd1 (tGnd1) (Heck *et al.*, 2010) or a misfolded mutant of carboxypeptidase Y impaired in import into the endoplasmic reticulum (ΔssCPY^*^) (Park *et al.*, 2007) tagged with the mCherry-sfGFP timer. Whole cell extracts were separated by SDS-PAGE and probed with antibodies against YFP and Zwf1 as loading control. Three major forms observed for each fusion are indicated as in Figure 2D. Accumulation of processed tFT fragments is reduced in strains lacking the Ubr1 and San1 E3 ubiquitin ligases, which mark tGnd1 and ΔssCPY* for proteasomal degradation (Heck *et al.*, 2010). E – Degradation of a temperature-sensitive mutant of firefly luciferase (LuciDM) (Gupta *et al.*, 2011) tagged with the mCherry-sfGFP timer. Ubr1 and San1 target LuciDM for proteasomal degradation. Whole cell extracts of samples collected at the indicated time points after blocking translation with cycloheximide were separated by SDS-PAGE and probed with antibodies against YFP and Zwf1 as loading control. Three major forms observed for each Ubi-X-mCherry-sfGFP fusion are indicated as in Figure 2D. F – Incomplete proteasomal degradation of sfGFP but not mCherry fusions. Whole cell extracts of strains expressing the indicating fusions were separated by SDS-PAGE and probed with antibodies against the HA or myc epitopes. Major forms are indicated as in Figure 2D, cross-reacting background bands are marked with asterisks.

#### 2.3. Supplemental Figure S3

**Figure S3.**
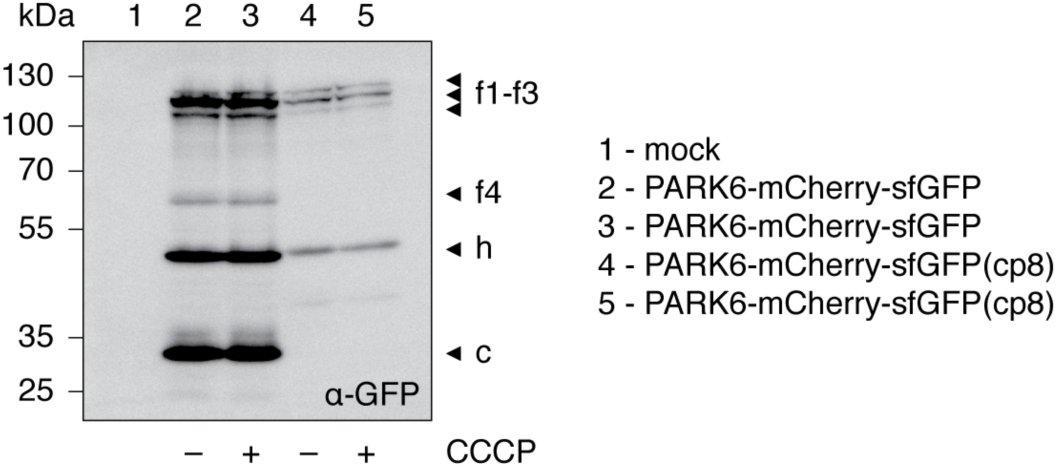
Comparison of mCherry-sfGFP and mCherry-sfGFP(cp8) timers in HEK293T cells. Immunoblot of HEK293T cells expressing PARK6-mCherry-sfGFP or PARK6-mCherry-sfGFP(cp8) with or without exposure to carbonyl cyanide *m*-chlorophenyl hydrazone (CCCP), which disrupts the mitochondrial membrane potential. Whole cell extracts were separated by SDS-PAGE and probed with an antibody against GFP. Major forms observed for each tFT fusion are indicated: full length fusions and PARK6 cleavage fragments generated by mitochondrial proteases (f1-f4) (Meissner *et al.*, 2011), a shorter mCherry^ΔN^ - greenFP product resulting from mCherry hydrolysis during cell extract preparation (h) (Gross *et al.*, 2000) and fast-migrating processed tFT fragments (c). Accumulation of processed tFT fragments does not depend on mitochondrial import of PARK6, as demonstrated by the insensitivity to CCCP, but is strongly reduced by the cp8 permutation of sfGFP.

#### 2.4. Supplemental Figure S4

**Figure S4.**
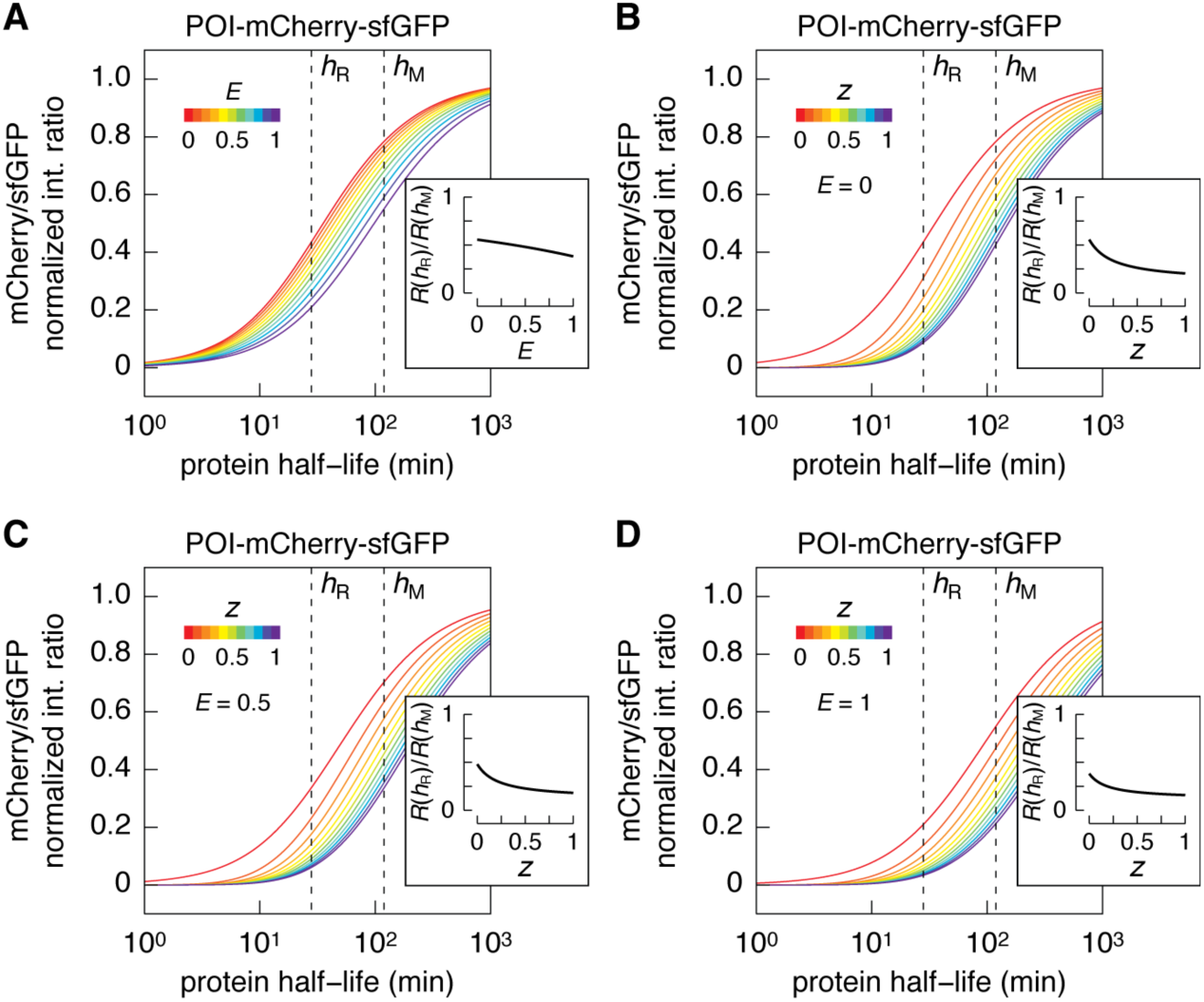
Theoretical analysis of tFTs with different FRET efficiencies between sfGFP and mCherry. A – Relationship between half-life of a protein of interest (POI) tagged at the C-terminus with mCherry-sfGFP and mCherry/sfGFP intensity ratio in steady state. mCherry/sfGFP intensity ratios were calculated as a function of protein degradation kinetics for a population doubling time of 90 min using published maturation parameters for mCherry and sfGFP (Khmelinskii *et al.*, 2012), a probability of incomplete degradation of sfGFP *z* = 0 and a FRET efficiency *E* between 0 and 1 (FRET from sfGFP to mCherry). Each curve is normalized to the mCherry/sfGFP intensity ratio of a non-degradable tFT fusion. Curves are color-coded according to *E* as indicated. Further details are provided as Supplemental Theory. Inset shows the comparison of mCherry/sfGFP intensity ratios (*R*) for two protein half-lives – *h*_R_ = 28 min (half-life of R-mCherry-sfGFP) and *h*_M_ = 119 min (half-life of M-mCherry-sfGFP) (Figure S2B) – as a function of FRET efficiency *E*. B-D – Relationship between half-life of a protein of interest (POI) tagged at the C-terminus with mCherry-sfGFP and mCherry/sfGFP intensity ratio in steady state. mCherry/sfGFP intensity ratios were calculated as a function of protein degradation kinetics for a population doubling time of 90 min using published maturation parameters for mCherry and sfGFP (Khmelinskii *et al.*, 2012), a degradation rate constant of free sfGFP *k*_2_ = 0, a FRET efficiency *E* of 0 (B), 0.5 (C) and 1 (D) (FRET from sfGFP to mCherry), and a probability *z* of incomplete degradation of sfGFP between 0 and 1. Each curve is normalized to the mCherry/sfGFP intensity ratio of a non-degradable tFT fusion. Curves are color-coded according to *z* as indicated. Further details are provided as Supplemental Theory. Insets show the comparison of mCherry/sfGFP intensity ratios (*R*) for two protein half-lives – h_R_ = 28 min (half-life of R-mCherry-sfGFP) and *h*_M_ = 119 min (half-life of M-mCherry-sfGFP) (Figure S2B) – as a function of *z*. Panel B is identical to Figure 6A and is shown for ease of comparison.

#### 2.5. Supplemental Figure S5

**Figure S5.**
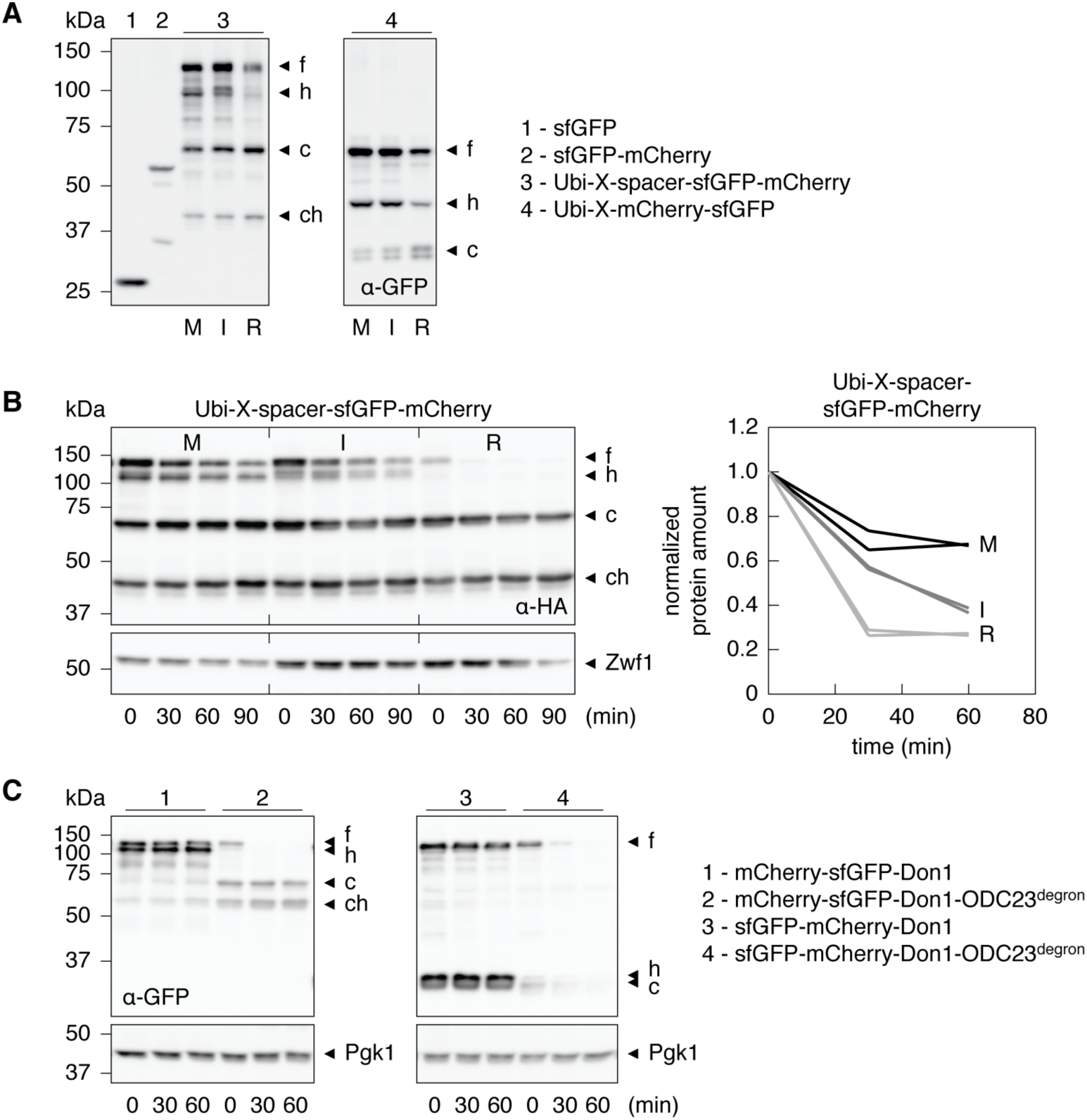
Degradation of tFT fusions with different ordering of fluorescent proteins in the timer. A – Whole cell extracts of strains expressing the indicated constructs were separated by SDS-PAGE and probed with antibodies against GFP. The residue X in each Ubi-X-tFT fusion is specified in the immunoblots. Four major forms observed for each Ubi-X-spacer-sfGFP-mCherry fusion are indicated: a full-length X-spacer-sfGFP-mCherry form (f), a shorter X-spacer-sfGFP-mCherry^ΔC^ product resulting from mCherry hydrolysis during cell extract preparation (h), fast-migrating processed tFT fragments (c) and corresponding hydrolysis products (ch). Three major forms observed for each Ubi-X-mCherry-sfGFP fusion are indicated as in Figure 2D. B – Degradation of Ubi-X-spacer-sfGFP-mCherry fusions after blocking translation with cycloheximide. Whole cell extracts of samples collected at the indicated time points were separated by SDS-PAGE and probed with antibodies against the HA epitope and Zwf1 as loading control. Four major forms observed for each fusion are indicated as in A. Quantification of two replicates of the cycloheximide chase is shown in the right panel. The combined amount of the (f) and (h) forms was normalized to the starting amount for each replicate. The residue X in each fusion is specified in the plot. C – Degradation of tFT-Don1 fusions, with and without a C-terminal ODC23 degron, after blocking translation with cycloheximide. Whole cell extracts of samples collected at the indicated time points were separated by SDS-PAGE and probed with antibodies against GFP and Pgk1 as loading control. Major forms observed for each tFT-Don1 fusion are indicated as in A.

#### 2.6. Supplemental Figure S6

**Figure S6.**
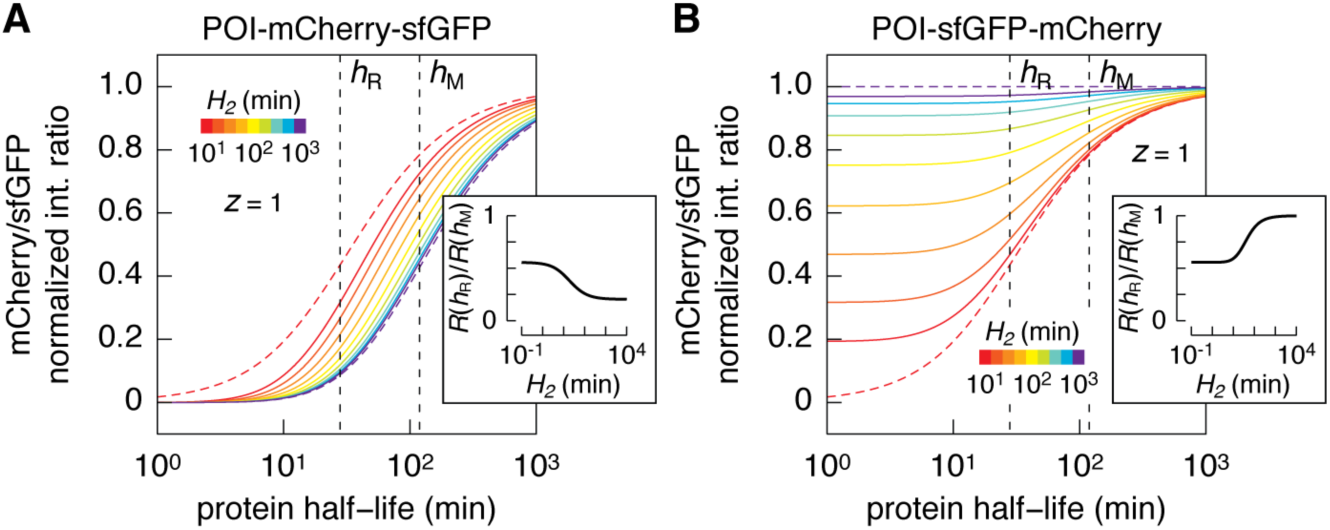
Theoretical analysis of tFTs with different degradation rates of processed tFT fragments. A, B – Relationship between half-life of a protein of interest (POI) tagged at the C-terminus with a tFT and mCherry/sfGFP intensity ratio in steady state. Incomplete degradation produces free sfGFP in A and free sfGFP-mCherry in B. mCherry/sfGFP intensity ratios were calculated as a function of protein degradation kinetics for a population doubling time of 90 min using published maturation parameters for mCherry and sfGFP (Khmelinskii *et al.*, 2012), a probability of incomplete degradation of tFT fusions *z* = 1 and a degradation rate constant of processed tFT fragments *k*_2_ between 0 and infinity. Note that degradation rate constant *k*_2_ is related to half-life *H*_2_ as *k*_2_ = ln(2)/*H*_2_. Each curve is normalized to the mCherry/sfGFP intensity ratio of a non-degradable tFT fusion. Full curves are color-coded according to *H*_2_ as indicated. Dashed curves correspond to *k*_2_ = 0 (purple, shown in Figure 6, A and B) and *k*_2_ = < (red). Further details are provided as Supplemental Theory. Insets show the comparison of mCherry/sfGFP intensity ratios (*R*) for two protein half-lives – h_R_ = 28 min (half-life of R-mCherry-sfGFP) and *h*_M_ = 119 min (half-life of M-mCherry-sfGFP) (Figure S2B) – as a function of *H*_2_.

### 3. Supplemental Tables

#### 3.1. Supplemental Table S1 – Strains used in this study

| Strain | Background | Genotype | Figure | Reference |
| --- | --- | --- | --- | --- |
| FY1679 | S288c | MATa/MATalpha ura3-52/ura3-52 leu2Δ1/LEU2<br>his3Δ200/HIS3 trp1Δ63/TRP1 GAL2/GAL2 |  | (Winston *et al.*, 1995) |
| ESM356-1 | FY1679 | MATa ura3-52 leu2Δ1 his3Δ200 trp1Δ63 |  | Elmar Schiebel |
| Y8205 | S288c | MATalpha his3Δ1 leu2Δ0 ura3Δ0 met15Δ0<br>can1Δ::P<sub>STE2</sub>-his5 lyp1Δ::P<sub>STE3</sub>-LEU2 |  | (Tong and Boone, 2007) |
| yJLM11-1 | Y8205 | URA3 |  |  |
| AK1093 | ESM356-1 | PMA1-mCherry-sfGFP-kanMX | 1 |  |
| yMaM659 | ESM356-1 | PMA1-mCherry-Clover-kanMX | 1 |  |
| yMaM647 | ESM356-1 | PMA1-mCherry-myeGFP-kanMX | 1 |  |
| yMaM651 | ESM356-1 | PMA1-mCherry-GFPm<sup>A206R</sup>-kanMX | 1 |  |
| yMaM655 | ESM356-1 | PMA1-mCherry-mNeonGreen-HA-kanMX | 1 |  |
| yMaM982 | ESM356-1 | PMA1-mCherry-sfGFP(cp3) | 1 |  |
| yMaM983 | ESM356-1 | PMA1-mCherry-sfGFP(cp7) | 1 |  |
| yMaM984 | ESM356-1 | PMA1-mCherry-sfGFP(cp8) | 1 |  |
| yMaM985 | ESM356-1 | PMA1-mCherry-sfGFP(cp9) | 1 |  |
| yMaM683 | ESM356-1 | PMA1-sfGFP-mCherry-kanMX | 1 |  |
| yMaM38 | ESM356-1 | pRS415-P<sub>GPD</sub>-Ubi-M-mCherry-sfGFP | 2-6,S5 |  |
| yMaM44 | ESM356-1 | pRS415-P<sub>GPD</sub>-Ubi-R-mCherry-sfGFP | 2-6,S5 |  |
| yMaM620 | ESM356-1 | pRS415-P<sub>GPD</sub>-Ubi-M-mCherry-myeGFP | 2 |  |
| yMaM624 | ESM356-1 | pRS415-P<sub>GPD</sub>-Ubi-R-mCherry-myeGFP | 2 |  |
| yMaM621 | ESM356-1 | pRS415-P<sub>GPD</sub>-Ubi-M-mCherry-GFPm<sup>A206R</sup> | 2 |  |
| yMaM622 | ESM356-1 | pRS415-P<sub>GPD</sub>-Ubi-R-mCherry-GFPm<sup>A206R</sup> | 2 |  |
| yMaM616 | ESM356-1 | pRS415-P<sub>GPD</sub>-Ubi-M-mCherry-Clover | 2,4 |  |
| yMaM619 | ESM356-1 | pRS415-P<sub>GPD</sub>-Ubi-R-mCherry-Clover | 2,4 |  |
| yMaM625 | ESM356-1 | pRS415-P<sub>GPD</sub>-Ubi-M-mCherry-mNeonGreen-HA | 2 |  |
| yMaM623 | ESM356-1 | pRS415-P<sub>GPD</sub>-Ubi-R-mCherry-mNeonGreen-HA | 2 |  |
| yMaM32 | ESM356-1 | pRS415-P<sub>GPD</sub>-sfGFP | 2-4,S1,S5 |  |
| yMaM699 | ESM356-1 | pRS415-P<sub>GPD</sub>-mCherry-sfGFP | 3 |  |
| yMaM666 | ESM356-1 | pRS415-P<sub>GPD</sub>-Ubi-T-mCherry-sfGFP | 3,6 |  |
| yMaM35 | ESM356-1 | pRS415-P<sub>GPD</sub>-Ubi-I-mCherry-sfGFP | 3,4,6,S5 |  |
| yMaM108 | ESM356-1 | pRS415-P<sub>GPD</sub>-Ubi-Y-mCherry-sfGFP | 3,6 |  |
| yMaM41 | ESM356-1 | pRS415-P<sub>GPD</sub>-Ubi-F-mCherry-sfGFP | 3,6 |  |
| yMaM597 | ESM356-1 | pRS415-P<sub>GPD</sub>-Ubi-M-mCherry-LDGAG-sfGFP | 4 |  |
| yMaM598 | ESM356-1 | pRS415-P<sub>GPD</sub>-Ubi-I-mCherry-LDGAG-sfGFP | 4 |  |
| yMaM599 | ESM356-1 | pRS415-P<sub>GPD</sub>-Ubi-R-mCherry-LDGAG-sfGFP | 4 |  |
| yMaM600 | ESM356-1 | pRS415-P<sub>GPD</sub>-Ubi-M-mCherry-LDGS-sfGFP | 4 |  |
| yMaM601 | ESM356-1 | pRS415-P<sub>GPD</sub>-Ubi-I-mCherry-LDGS-sfGFP | 4 |  |
| yMaM602 | ESM356-1 | pRS415-P<sub>GPD</sub>-Ubi-R-mCherry-LDGS-sfGFP | 4 |  |
| yMaM711 | ESM356-1 | pRS415-P<sub>GPD</sub>-Ubi-M-mCherry-sfGFP<sup>L64F</sup> | 4 |  |
| yMaM712 | ESM356-1 | pRS415-P<sub>GPD</sub>-Ubi-R-mCherry-sfGFP<sup>L64F</sup> | 4 |  |
| yMaM715 | ESM356-1 | pRS415-P<sub>GPD</sub>-Ubi-M-mCherry-sfGFP<sup>5M</sup> | 4 |  |
| yMaM716 | ESM356-1 | pRS415-P<sub>GPD</sub>-Ubi-R-mCherry-sfGFP<sup>5M</sup> | 4 |  |
| yMaM717 | ESM356-1 | pRS415-P<sub>GPD</sub>-Ubi-M-mCherry-Clover<sup>F64L</sup> | 4 |  |
| yMaM718 | ESM356-1 | pRS415-P<sub>GPD</sub>-Ubi-R-mCherry-Clover<sup>F64L</sup> | 4 |  |
| yMaM968 | ESM356-1 | pRS415-P<sub>GPD</sub>-Ubi-M-mCherry-sfGFP<sup>cp3</sup> | 5 |  |
| yMaM969 | ESM356-1 | pRS415-P<sub>GPD</sub>-Ubi-M-mCherry-sfGFP<sup>cp7</sup> | 5 |  |
| yMaM970 | ESM356-1 | pRS415-P<sub>GPD</sub>-Ubi-M-mCherry-sfGFP<sup>cp8</sup> | 5 |  |
| yMaM971 | ESM356-1 | pRS415-P<sub>GPD</sub>-Ubi-M-mCherry-sfGFP<sup>cp9</sup> | 5 |  |
| yMaM972 | ESM356-1 | pRS415-P<sub>GPD</sub>-Ubi-M-mCherry-sfGFP<sup>cp12</sup> | 5 |  |
| yMaM973 | ESM356-1 | pRS415-P<sub>GPD</sub>-Ubi-M-mCherry-sfGFP<sup>cp13</sup> | 5 |  |
| yMaM974 | ESM356-1 | pRS415-P<sub>GPD</sub>-Ubi-R-mCherry-sfGFP<sup>cp3</sup> | 5 |  |
| yMaM975 | ESM356-1 | pRS415-P<sub>GPD</sub>-Ubi-R-mCherry-sfGFP<sup>cp7</sup> | 5 |  |
| yMaM976 | ESM356-1 | pRS415-P<sub>GPD</sub>-Ubi-R-mCherry-sfGFP<sup>cp8</sup> | 5 |  |
| yMaM977 | ESM356-1 | pRS415-P<sub>GPD</sub>-Ubi-R-mCherry-sfGFP<sup>cp9</sup> | 5 |  |
| yMaM978 | ESM356-1 | pRS415-P<sub>GPD</sub>-Ubi-R-mCherry-sfGFP<sup>cp12</sup> | 5 |  |
| yMaM979 | ESM356-1 | pRS415-P<sub>GPD</sub>-Ubi-R-mCherry-sfGFP<sup>cp13</sup> | 5 |  |
| yMaM26 | ESM356-1 | pRS415-P<sub>GPD</sub> | 6,S1,S2 |  |
| yMaM723 | ESM356-1 | pRS415-P<sub>GPD</sub>-Ubi-R-Don1-HA-sfGFP-mCherry | 6,S5 |  |
| yMaM728 | ESM356-1 | pRS415-P<sub>GPD</sub>-Ubi-F-Don1-HA-sfGFP-mCherry | 6 |  |
| yMaM740 | ESM356-1 | pRS415-P<sub>GPD</sub>-Ubi-Y-Don1-HA-sfGFP-mCherry | 6 |  |
| yMaM722 | ESM356-1 | pRS415-P<sub>GPD</sub>-Ubi-I-Don1-HA-sfGFP-mCherry | 6,S5 |  |
| yMaM721 | ESM356-1 | pRS415-P<sub>GPD</sub>-Ubi-M-Don1-HA-sfGFP-mCherry | 6,S5 |  |
| yMaM737 | ESM356-1 | pRS415-P<sub>GPD</sub>-Ubi-T-Don1-HA-sfGFP-mCherry | 6 |  |
| yMaM958 | ESM356-1 | pRS415-P<sub>GPD</sub>-mCherry-sfGFP-Don1 | 6,S5 |  |
| yMaM959 | ESM356-1 | pRS415-P<sub>GPD</sub>-mCherry-sfGFP-Don1-ODC23 | 6,S5 |  |
| yMaM960 | ESM356-1 | pRS415-P<sub>GPD</sub>-sfGFP-mCherry-Don1 | 6,S5 |  |
| yMaM961 | ESM356-1 | pRS415-P<sub>GPD</sub>-sfGFP-mCherry-Don1-ODC23 | 6,S5 |  |
| AK1169 | ESM356-1 | ura3-52::P<sub>GDP</sub>-Ubi-M-mCherry-sfGFP::kanMX | S2 | (Khmelniskii *et al.*, 2012) |
| AK1170 | ESM356-1 | ura3-52::P<sub>GDP</sub>-Ubi-R-mCherry-sfGFP::kanMX | S2 | (Khmelniskii *et al.*, 2012) |
| yMaM67 | ESM356-1 | pRS415-P<sub>GAL1</sub>-Ubi-R-mCherry-sfGFP | S2 |  |
| AK897 | ESM356-1 | pdr5Δ::kanMX | S2 |  |
| yMaM957 | ESM356-1 | pdr5Δ::kanMX pRS415-P<sub>GAL1</sub>-Ubi-R-mCherry-sfGFP | S2 |  |
| CHY108 | yJLM11-1 | ura3Δ::P<sub>TEF</sub>-tGnd1-mCherry-sfGFP::natNT2 | S2 |  |
| CHY109 | yJLM11-1 | ura3Δ::P<sub>TEF</sub>-tGnd1-mCherry-sfGFP::natNT2<br>ubr1Δ::hphNT1 | S2 |  |
| CHY110 | yJLM11-1 | ura3Δ::P<sub>TEF</sub>-tGnd1-mCherry-sfGFP::natNT2<br>san1Δ::kanMX4 | S2 |  |
| CHY111 | yJLM11-1 | ura3Δ::P<sub>TEF</sub>-tGnd1-mCherry-sfGFP::natNT2<br>ubr1Δ::hphNT1 san1Δ::kanMX4 | S2 |  |
| CHY112 | yJLM11-1 | ura3Δ::P<sub>TEF</sub>-ΔssCPY*-mCherry-sfGFP::natNT2 | S2 |  |
| CHY113 | yJLM11-1 | ura3Δ::P<sub>TEF</sub>-ΔssCPY*-mCherry-sfGFP::natNT2<br>ubr1Δ::hphNT1 | S2 |  |
| CHY114 | yJLM11-1 | ura3Δ::P<sub>TEF</sub>-ΔssCPY*-mCherry-sfGFP::natNT2<br>san1Δ::kanMX4 | S2 |  |
| CHY115 | yJLM11-1 | ura3Δ::P<sub>TEF</sub>-ΔssCPY*-mCherry-sfGFP::natNT2<br>ubr1Δ::hphNT1 san1Δ::kanMX4 | S2 |  |
| CHY116 | yJLM11-1 | ura3Δ::P<sub>TEF</sub>-LuciDM-mCherry-sfGFP::natNT2 | S2 |  |
| CHY117 | yJLM11-1 | ura3Δ::P<sub>TEF</sub>-LuciDM-mCherry-sfGFP::natNT2<br>ubr1Δ::hphNT1 | S2 |  |
| CHY118 | yJLM11-1 | ura3Δ::P<sub>TEF</sub>-LuciDM-mCherry-sfGFP::natNT2<br>san1Δ::kanMX4 | S2 |  |
| CHY119 | yJLM11-1 | ura3Δ::P<sub>TEF</sub>-LuciDM-mCherry-sfGFP::natNT2<br>ubr1Δ::hphNT1 san1Δ::kanMX4 | S2 |  |
| yMaM1072 | ESM356-1 | pRS415-P<sub>GDP</sub>-Ubi-R-Don1-HA-sfGFP-myc | S2 |  |
| yMaM1070 | ESM356-1 | pRS415-P<sub>GDP</sub>-Ubi-R-Don1-HA-mCherry-myc | S2 |  |
| yMaM700 | ESM356-1 | pRS415-P<sub>GDP</sub>-sfGFP-mCherry | S5 |  |

#### 3.2. Supplemental Table S2 – Plasmids used in this study

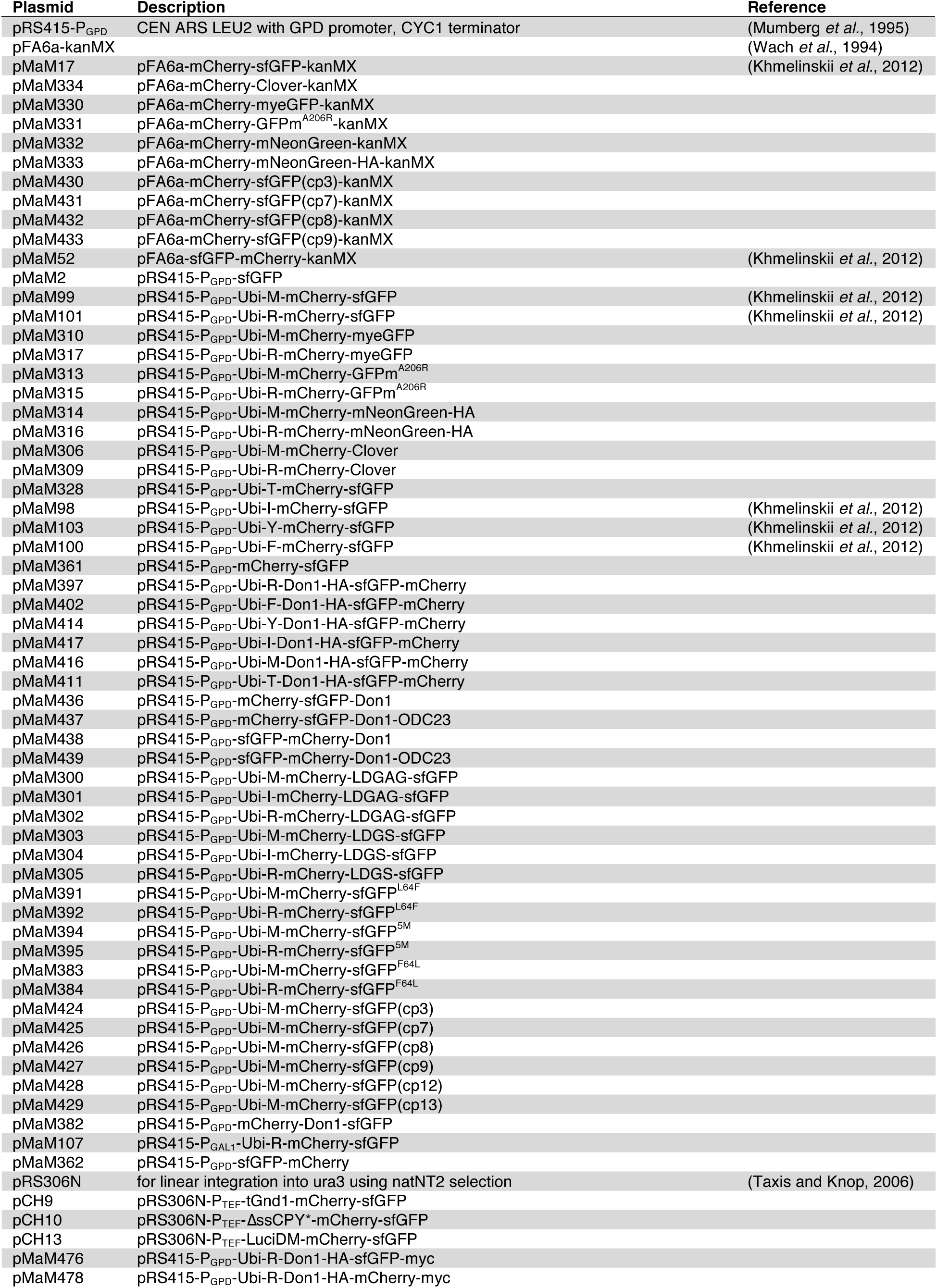

| Plasmid | Description | Reference |
| --- | --- | --- |
| pRS415-P<sub>GPD</sub> | CEN ARS LEU2 with GPD promoter, CYC1 terminator | (Mumberg *et al.*, 1995) |
| pFA6a-kanMX |  | (Wach *et al.*, 1994) |
| pMaM17 | pFA6a-mCherry-sfGFP-kanMX | (Khmelinskii *et al.*, 2012) |
| pMaM334 | pFA6a-mCherry-Clover-kanMX |  |
| pMaM330 | pFA6a-mCherry-myeGFP-kanMX |  |
| pMaM331 | pFA6a-mCherry-GFPm<sup>A206R</sup>-kanMX |  |
| pMaM332 | pFA6a-mCherry-mNeonGreen-kanMX |  |
| pMaM333 | pFA6a-mCherry-mNeonGreen-HA-kanMX |  |
| pMaM430 | pFA6a-mCherry-sfGFP(cp3)-kanMX |  |
| pMaM431 | pFA6a-mCherry-sfGFP(cp7)-kanMX |  |
| pMaM432 | pFA6a-mCherry-sfGFP(cp8)-kanMX |  |
| pMaM433 | pFA6a-mCherry-sfGFP(cp9)-kanMX |  |
| pMaM52 | pFA6a-sfGFP-mCherry-kanMX | (Khmelinskii *et al.*, 2012) |
| pMaM2 | pRS415-P<sub>GPD</sub>-sfGFP |  |
| pMaM99 | pRS415-P<sub>GPD</sub>-Ubi-M-mCherry-sfGFP | (Khmelinskii *et al.*, 2012) |
| pMaM101 | pRS415-P<sub>GPD</sub>-Ubi-R-mCherry-sfGFP | (Khmelinskii *et al.*, 2012) |
| pMaM310 | pRS415-P<sub>GPD</sub>-Ubi-M-mCherry-myeGFP |  |
| pMaM317 | pRS415-P<sub>GPD</sub>-Ubi-R-mCherry-myeGFP |  |
| pMaM313 | pRS415-P<sub>GPD</sub>-Ubi-M-mCherry-GFPm<sup>A206R</sup> |  |
| pMaM315 | pRS415-P<sub>GPD</sub>-Ubi-R-mCherry-GFPm<sup>A206R</sup> |  |
| pMaM314 | pRS415-P<sub>GPD</sub>-Ubi-M-mCherry-mNeonGreen-HA |  |
| pMaM316 | pRS415-P<sub>GPD</sub>-Ubi-R-mCherry-mNeonGreen-HA |  |
| pMaM306 | pRS415-P<sub>GPD</sub>-Ubi-M-mCherry-Clover |  |
| pMaM309 | pRS415-P<sub>GPD</sub>-Ubi-R-mCherry-Clover |  |
| pMaM328 | pRS415-P<sub>GPD</sub>-Ubi-T-mCherry-sfGFP |  |
| pMaM98 | pRS415-P<sub>GPD</sub>-Ubi-I-mCherry-sfGFP | (Khmelinskii *et al.*, 2012) |
| pMaM103 | pRS415-P<sub>GPD</sub>-Ubi-Y-mCherry-sfGFP | (Khmelinskii *et al.*, 2012) |
| pMaM100 | pRS415-P<sub>GPD</sub>-Ubi-F-mCherry-sfGFP | (Khmelinskii *et al.*, 2012) |
| pMaM361 | pRS415-P<sub>GPD</sub>-mCherry-sfGFP |  |
| pMaM397 | pRS415-P<sub>GPD</sub>-Ubi-R-Don1-HA-sfGFP-mCherry |  |
| pMaM402 | pRS415-P<sub>GPD</sub>-Ubi-F-Don1-HA-sfGFP-mCherry |  |
| pMaM414 | pRS415-P<sub>GPD</sub>-Ubi-Y-Don1-HA-sfGFP-mCherry |  |
| pMaM417 | pRS415-P<sub>GPD</sub>-Ubi-I-Don1-HA-sfGFP-mCherry |  |
| pMaM416 | pRS415-P<sub>GPD</sub>-Ubi-M-Don1-HA-sfGFP-mCherry |  |
| pMaM411 | pRS415-P<sub>GPD</sub>-Ubi-T-Don1-HA-sfGFP-mCherry |  |
| pMaM436 | pRS415-P<sub>GPD</sub>-mCherry-sfGFP-Don1 |  |
| pMaM437 | pRS415-P<sub>GPD</sub>-mCherry-sfGFP-Don1-ODC23 |  |
| pMaM438 | pRS415-P<sub>GPD</sub>-sfGFP-mCherry-Don1 |  |
| pMaM439 | pRS415-P<sub>GPD</sub>-sfGFP-mCherry-Don1-ODC23 |  |
| pMaM300 | pRS415-P<sub>GPD</sub>-Ubi-M-mCherry-LDGAG-sfGFP |  |
| pMaM301 | pRS415-P<sub>GPD</sub>-Ubi-I-mCherry-LDGAG-sfGFP |  |
| pMaM302 | pRS415-P<sub>GPD</sub>-Ubi-R-mCherry-LDGAG-sfGFP |  |
| pMaM303 | pRS415-P<sub>GPD</sub>-Ubi-M-mCherry-LDGS-sfGFP |  |
| pMaM304 | pRS415-P<sub>GPD</sub>-Ubi-I-mCherry-LDGS-sfGFP |  |
| pMaM305 | pRS415-P<sub>GPD</sub>-Ubi-R-mCherry-LDGS-sfGFP |  |
| pMaM391 | pRS415-P<sub>GPD</sub>-Ubi-M-mCherry-sfGFP<sup>L64F</sup> |  |
| pMaM392 | pRS415-P<sub>GPD</sub>-Ubi-R-mCherry-sfGFP<sup>L64F</sup> |  |
| pMaM394 | pRS415-P<sub>GPD</sub>-Ubi-M-mCherry-sfGFP<sup>5M</sup> |  |
| pMaM395 | pRS415-P<sub>GPD</sub>-Ubi-R-mCherry-sfGFP<sup>5M</sup> |  |
| pMaM383 | pRS415-P<sub>GPD</sub>-Ubi-M-mCherry-sfGFP<sup>F64L</sup> |  |
| pMaM384 | pRS415-P<sub>GPD</sub>-Ubi-R-mCherry-sfGFP<sup>F64L</sup> |  |
| pMaM424 | pRS415-P<sub>GPD</sub>-Ubi-M-mCherry-sfGFP(cp3) |  |
| pMaM425 | pRS415-P<sub>GPD</sub>-Ubi-M-mCherry-sfGFP(cp7) |  |
| pMaM426 | pRS415-P<sub>GPD</sub>-Ubi-M-mCherry-sfGFP(cp8) |  |
| pMaM427 | pRS415-P<sub>GPD</sub>-Ubi-M-mCherry-sfGFP(cp9) |  |
| pMaM428 | pRS415-P<sub>GPD</sub>-Ubi-M-mCherry-sfGFP(cp12) |  |
| pMaM429 | pRS415-P<sub>GPD</sub>-Ubi-M-mCherry-sfGFP(cp13) |  |
| pMaM382 | pRS415-P<sub>GPD</sub>-mCherry-Don1-sfGFP |  |
| pMaM107 | pRS415-P<sub>GAL1</sub>-Ubi-R-mCherry-sfGFP |  |
| pMaM362 | pRS415-P<sub>GPD</sub>-sfGFP-mCherry |  |
| pRS306N | for linear integration into ura3 using natNT2 selection | (Taxis and Knop, 2006) |
| pCH9 | pRS306N-P<sub>TEF</sub>-tGnd1-mCherry-sfGFP |  |
| pCH10 | pRS306N-P<sub>TEF</sub>-ΔssCPY*-mCherry-sfGFP |  |
| pCH13 | pRS306N-P<sub>TEF</sub>-LuciDM-mCherry-sfGFP |  |
| pMaM476 | pRS415-P<sub>GPD</sub>-Ubi-R-Don1-HA-sfGFP-myc |  |
| pMaM478 | pRS415-P<sub>GPD</sub>-Ubi-R-Don1-HA-mCherry-myc |  |

